# Gata2, Nkx2-2 and Skor2 form a transcription factor network regulating development of a midbrain GABAergic neuron subtype with characteristics of REM sleep regulatory neurons

**DOI:** 10.1101/2022.01.05.473755

**Authors:** Anna Kirjavainen, Parul Singh, Laura Lahti, Patricia Seja, Zoltan Lelkes, Aki Makkonen, Sami Kilpinen, Yuichi Ono, Marjo Salminen, Teemu Aitta-Aho, Tarja Stenberg, Svetlana Molchanova, Kaia Achim, Juha Partanen

**Affiliations:** Molecular and Integrative Biosciences Research Programme, Faculty of Biological and Environmental Sciences, P. O. Box 56, FIN00014-University of Helsinki, Helsinki, Finland; Department of Physiology, P.O. Box 63, FIN00014-University of Helsinki, Helsinki, Finland; Department of Pharmacology, P.O. Box 63, FIN00014-University of Helsinki, Helsinki, Finland; Department of Physiology, Faculty of Medicine, University of Szeged, Szeged, Hungary; Integrated Cell Biology, KAN Research Institute, Inc., 6-8-2 Minatojima-Minamimachi, Chuo-ku, Kobe, Hyogo 650-0047, Japan; Department of Veterinary Biosciences, P. O. Box 66, FIN00014-University of Helsinki, Helsinki, Finland

**Keywords:** midbrain reticular formation, deep mesencephalic nucleus, periaqueductal gray, neurogenesis, transcription factor, Skor2, Nkx2-2, Gata2, REM sleep, mouse, rat

## Abstract

The midbrain reticular formation is a mosaic of diverse GABAergic and glutamatergic neurons that have been associated with a variety of functions, including the regulation of sleep. However the molecular characteristics and development of the midbrain reticular formation neurons are poorly understood. As the transcription factor Gata2 is required for the development of all GABAergic neurons derived from the embryonic mouse midbrain, we hypothesized that the genes expressed downstream of Gata2 could contribute to the diversification of GABAergic neuron subtypes in this brain region. Here, we show that Gata2 is indeed required for the expression of several lineage-specific transcription factors in post-mitotic midbrain GABAergic neuron precursors. These include a homeodomain transcription factor Nkx2-2 and a SKI family transcriptional repressor Skor2, which are co-expressed in a restricted group of GABAergic precursors in the midbrain reticular formation. Both Gata2, and Nkx2-2 function is required for the expression of *Skor2* in GABAergic precursors. In the adult mouse as well as rat midbrain, the *Nkx2-2* and *Skor2* expressing GABAergic neurons locate at the boundary of the ventrolateral periaqueductal gray and the midbrain reticular formation, an area shown to contain REM-off neurons regulating REM sleep. In addition to the characteristic localization, the *Skor2* positive cells increase their activity upon REM sleep inhibition, send projections to a pontine region associated with sleep control and are responsive to orexins, consistent with the known properties of the midbrain REM-off neurons.

## Introduction

Gamma-aminobutyric acid (GABA) is the main inhibitory neurotransmitter in the mature brain and neurons using GABA as their principal transmitter (GABAergic neurons) are widely distributed throughout the central nervous system. In the midbrain, abundant GABAergic neurons are located in the midbrain reticular formation (MRF; also known as the deep mesencephalic nucleus) and the periaqueductal gray (PAG). Very little is known about the subtype-specific features of the GABAergic neurons in the MRF and PAG. This is a major obstacle for understanding the MRF and PAG associated brain functions, including regulation of defensive behavior, nociception, and sleep (Keay and Bandler, 2001; Luppi et al., 2017). A region at the boundary of the dorsomedial MRF (dMRF) and ventrolateral PAG (vlPAG) contains GABAergic neurons implicated in the regulation of rapid eye movement (REM) sleep. The GABAergic neurons in the dMRF/vlPAG are activated by REM sleep deprivation, project to other brainstem regions regulating REM sleep, and appear to modulate the sleep pattern (Luppi et al., 2017; Saper et al., 2010; Scammell et al., 2017; Weber and Dan, 2016). However, despite their functional importance, limited information is available on the development and differentiation of the MRF and PAG GABAergic neurons, and the subtype-specific molecular features of these cells.

In the embryonic midbrain, the development of GABAergic neuron precursors depends on a zinc-finger transcription factor (TF) Gata2. The expression of Gata2 is activated in midbrain precursors immediately upon their cell cycle exit, and Gata2 drives the GABAergic differentiation over alternative glutamatergic fates (Kala et al., 2009). Gata2 appears to function together with the Tal-family TFs, in particular Tal2, to direct GABAergic neurogenesis in the midbrain (Achim et al., 2013). The embryonic midbrain can be divided into dorso-ventral progenitor domains differing in their gene expression (m1-m7) (Kala et al., 2009; Nakatani et al., 2007). Of these, the domains m1-m3, ventral m4 and m5 give rise to post-mitotic GABAergic neuron precursors. While Gata2 and Tal2 are required for the differentiation of all the midbrain GABAergic precursors, the contribution of the precursor subtypes to brain nuclei, and the gene regulatory circuits guiding the subtype diversification, are unknown.

Here, we identify Gata2 dependent TFs marking midbrain GABAergic precursors and their subtypes. Of these, we focus on a restricted midbrain GABAergic neuron subtype defined by the coexpression of the homeodomain TF Nkx2-2 and the SKI family TF Skor2/Corl2. We show that in these cells, both Gata2 and Nkx2-2 are required for the expression of *Skor2*. We demonstrate that the Skor2 and Nkx2-2 co-expressing neuros have characteristics, such as anatomical location at the boundary of dMRF and vlPAG, activation by REM sleep deprivation, projection to pontine areas controlling REM sleep, and responsiveness to Orexin (hypocretin, Hcrt), suggesting that this group of midbrain GABAergic neurons is involved in REM-sleep regulation.

## Results

### Gene expression changes in the embryonic midbrain lacking Gata2 function

Gata2 operates high in the gene regulatory network guiding post-mitotic differentiation of GABAergic neuron precursors in the embryonic midbrain (Kala et al., 2009). To reveal the genes downstream of Gata2, we compared the gene expression in E12.5 control (*Ctrl*) and *En1^Cre/+^; Gata2^flox/flox^* (*Gata2^cko^*) mutant mouse midbrain (Fig. 1A, B). Using cDNA microarrays, we found 52 genes downregulated in the *Gata2^cko^* embryos, either in the ventral midbrain, dorsal midbrain, or both (logFC>1.5, Adjusted p-value<0.05, Supplementary Table S1; all up- and downregulated genes are listed in the Supplementary Table S2). We performed the qRT-PCR analyses of the expression of selected GABAergic neuron markers (*Gata2, Gad1, Slc32a1*), GATA-associated transcription factors (*Tal1, Zfpm1, Zfpm2*) and novel genes (*Ptchd4)*, across different magnitudes of fold changes. Overall, the fold change of each gene expression we observed between the *Ctrl* and *Gata2^cko^* in the qRT-PCR correlated well with the fold changes observed in the cDNA microarray comparisons (Supplementary Fig. S1). We validated the downregulation of selected genes also by mRNA in situ hybridization (ISH; Fig. 1C-O’; Supplementary Table S1).

**Figure 1.**
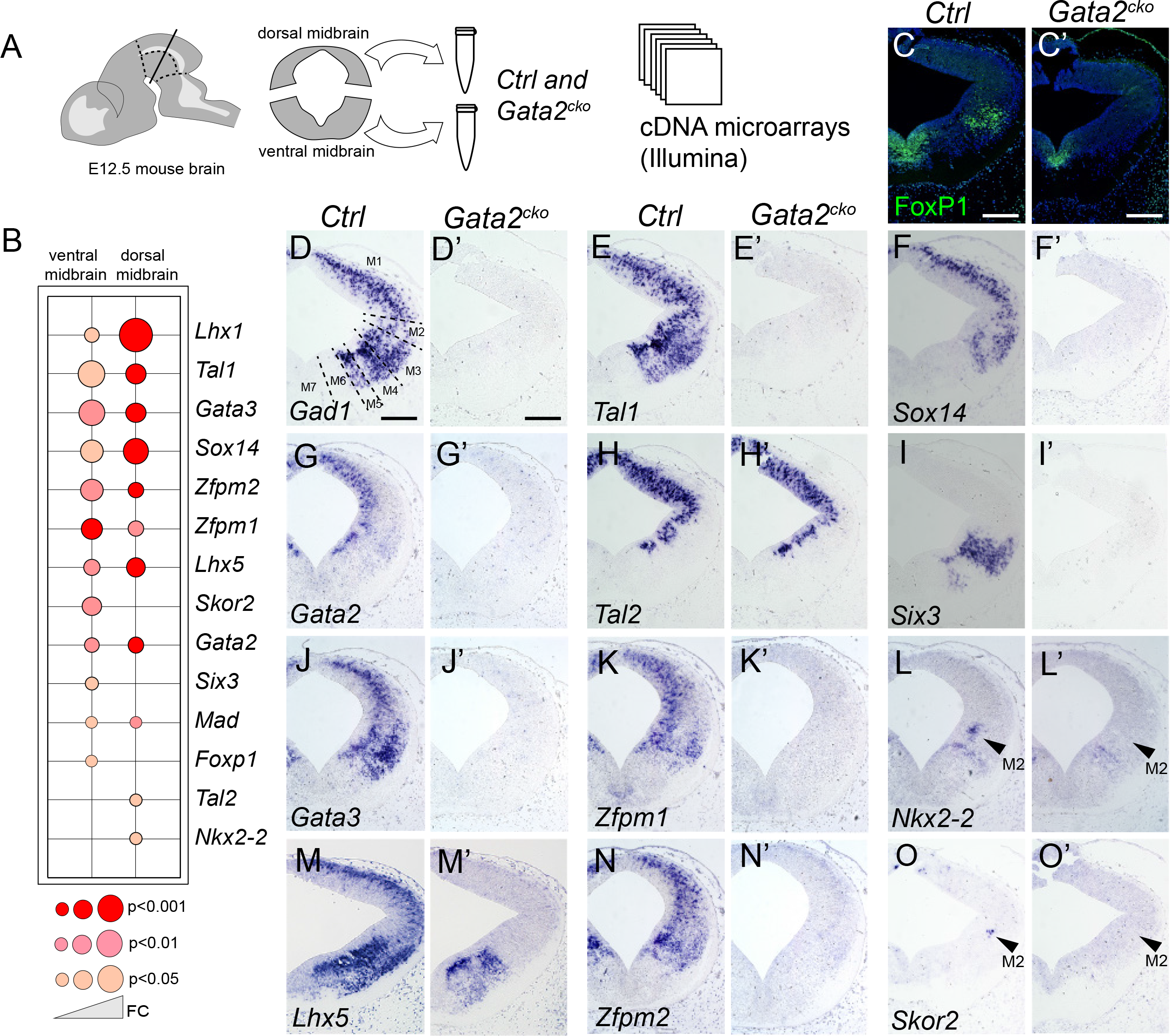
Gata2 regulated transcription factors in the developing mouse midbrain. A, Microarray sample collection and experimental design. The dorsal and ventral midbrain tissue from three E12.5 *Ctrl* and three E12.5 *Gata2^cko^* embryos was collected and analysed on cDNA microarrays. B, Downregulated transcription factor genes in the *Gata2^cko^* midbrain. Gene expression in *Ctrl* and *Gata2^cko^* embryos was compared separately for the ventral and dorsal midbrain samples. Dot size indicates fold change, the color indicates p-values. C-O’, IHC (C,C’) and ISH (D-O’) analysis of the expression of transcription factors identified as downregulated in the *Gata2^cko^* samples. Analyses were performed on coronal sections of E12.5 control (C-O) and *Gata2^cko^* (C’-O’) embryos. Arrowheads on L and O point to the location of a lateral cell population expressing *Nkx2-2* and *Skor2*, respectively. Scale bars, 100 um.

TF encoding genes were abundant among the ones down-regulated in the *Gata2^cko^* mutants (Supplementary Table S1; 14 out of 52 down-regulated genes encode TFs). The gene ontology (GO) term enrichment analysis listed ‘sequence specific DNA binding’ (p=1.06E-5), ‘positive regulation of transcription’ (p=4.63E-4), ‘E-box binding’ (p=0.009), ‘nervous system development’ (p=0.003), and ‘neuron differentiation’ (p=0.014) among top GO terms (Supplementary Table S3). As expected, the expression of several genes associated with GABAergic neuron functions (such as *Gad1, Gad2, Slc32a1*) were down-regulated in the *Gata2^cko^* mutants (Fig. 1 D-D’, Supplementary Table S1).

### Gata-associated TFs are broadly expressed in the midbrain GABAergic neuron precursors

To dissect the gene regulatory networks downstream of Gata2, we focused on the down-regulated TF genes (Fig. 1B). From those, *Gata3*, *Tal1*, *Tal2*, *Zfpm1* and *Zfpm2* have earlier been associated with the function of Gata2 or other Gata factors in different developmental contexts (Chlon and Crispino, 2012; Lahti et al., 2016; Morello et al., 2020; Tikker et al., 2020). ISH analysis showed that these putative Gata2 cofactors were broadly expressed in the E12.5 midbrain GABAergic neuron precursors (Fig. 1J, E-N). Consistent with the microarray profiling, and our earlier studies in the developing midbrain and diencephalon (Achim et al., 2013; Virolainen et al., 2012), *Tal1* expression was completely abolished in the *Gata2^cko^* (Fig. 1E-E’). In contrast, *Tal2* expression was only modestly downregulated in the dorsal midbrain sample and was still robustly expressed in the midbrain GABAergic precursors in the *Gata2^cko^* embryos (Fig. 1H-H’, Supplementary Table S1), supporting the hypothesis of independent activation of *Gata2* and *Tal2* expression and their position at the top of the gene regulatory hierarchy driving midbrain GABAergic neuron differentiation (Achim et al., 2013).

Both *Zfpm1* and *Zfpm2* encode for zinc-finger proteins associating with the Gata TF complex in other cell types (Chlon and Crispino, 2012). In E12.5 *Ctrl* midbrain, *Zfpm1* and *Zfpm2* transcripts were broadly expressed in the GABAergic neuron precursors, but undetectable in the *Gata2^cko^* midbrain (Fig. 1K-K’, N-N’). Thus, in the midbrain GABAergic precursors, the expression of several genes encoding for Gata-associated TFs requires Gata2.

Other Gata2 dependent TF genes expressed broadly in the midbrain GABAergic neuron precursors included the LIM-homeobox genes *Lhx1* and *Lhx5*. In contrast to the *Tal1/2* and *Zfpm1/2*, both *Lhx1* and *Lhx5* were also expressed in the glutamatergic neuron precursors in the m6 and m4 regions, where their expression was not affected by the loss of *Gata2* (Fig. 1M-M’ and data not shown).

### TFs downstream of Gata2 mark subtypes of mantle zone precursors in the embryonic midbrain

In contrast to the TFs expressed across all midbrain GABAergic neuron precursors, some Gata2 dependent TF genes had more restricted expression patterns. These included *FoxP1*, *Six3*, *Sox14*, *Nkx2-2* and *Skor2* (also known as *Fussel18* and *Corl2*). By ISH and IHC analyses at E12.5, we found *Sox14* expressing precursors primarily in the dorsal midbrain, in particular m1-m3 (Fig. 1F), consistent with earlier studies (Makrides et al., 2018). In turn, we detected precursors expressing *Six3* in the ventrolateral GABAergic domains m3 and m5, *FoxP1* in the m3, and Nkx2-2 in the m4 and m2 (Fig. 1I,C,L). Notably, *Skor2* showed the most restricted pattern of expression that partially resembled *Nkx2-2* expression in the m2 mantle zone (Fig. 1O). The expression of these TFs was lost in the mantle zone precursors in *Gata2^cko^* embryos (Fig. 1C-C’, F-F’, I-I’, L-L’, O-O’, Fig. 2C, F), except for *Nkx2-2,* which was abolished in the m2, but continued in glutamatergic precursors in the m4 domain, as well as proliferative progenitors in the ventral midbrain (Kala et al., 2009).

**Figure 2.**
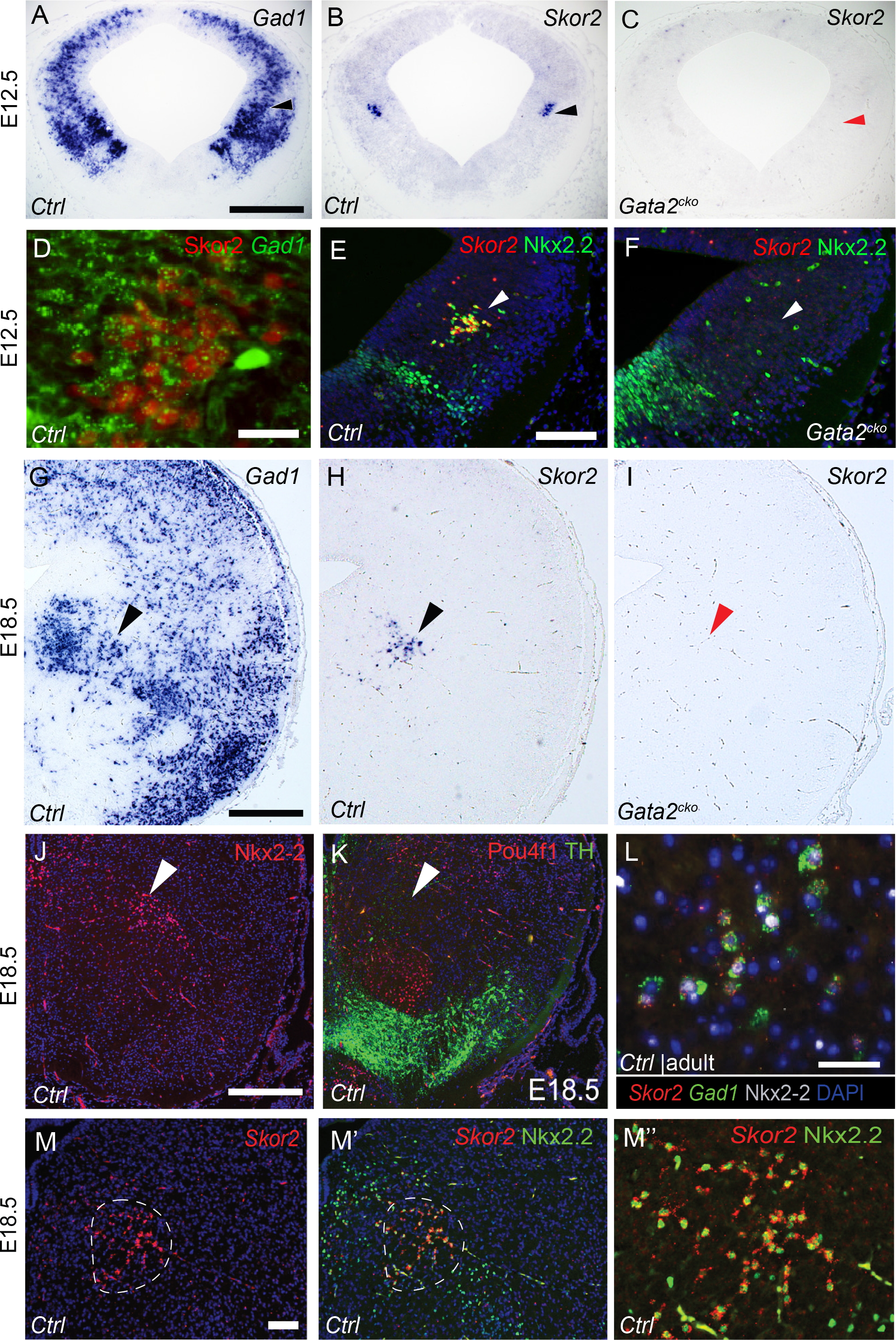
*Skor2* and *Nkx2-2* are co-expressed in a Gata2 dependent fashion in a subtype of lateral midbrain GABAergic precursors. A-B, Expression of *Gad1* (A) and *Skor2* (B) ISH on parallel coronal sections of an E12.5 *Ctrl* embryo. C, Loss of *Skor2* expression in an E12.5 *Gata2^cko^* embryo. D, Co-expression of Skor2 (IHC) and *Gad1* (ISH). Skor2 is detected in a subset of *Gad1* expressing cells in the lateral midbrain. E-F, Co-expression of Nkx2-2 (IHC) and *Skor2* (ISH) on E12.5 *Ctrl* and *Gata2^cko^* midbrain. The expression of both genes is lost in the *Gata2^cko^* lateral midbrain m2 domain (arrowhead). G, *Gad1* expression (ISH) on a coronal section of E18.5 *Ctrl* midbrain. H-I, *Skor2* expression (ISH) on a coronal section of E18.5 *Ctrl* and *Gata2^cko^* midbrain. J-K, Nkx2-2 (IHC), and TH and Pou4f1 (co-IHC) expression on adjacent coronal sections of E18.5 *Ctrl* embryo. L, Co-expression of Nkx2-2 (IHC), *Skor2* (ISH) and *Gad1* (ISH) in the adult mouse midbrain. Close-up image from the dMRF/vlPAG area corresponding to the E18.5 location indicated in panels J, K. M-M’’, Co-expression of *Skor2* (ISH) and Nkx2-2 (IHC) in the E18.5 mouse midbrain. M’’, close-up of the double labelled cells. The arrowheads point to the expected position of the *Skor2*^+^ cell population. Scale bars: 200 μm (A-C,H-L), 50 μm (D-G’’, M).

Varied, combinatorial expression pattern of the Gata2 target genes suggests that the development of diverse populations of midbrain GABAergic neurons entails unique Gata2 dependent TF networks in different post-mitotic GABAergic precursor populations.

### Co-expression of *Skor2* and *Nkx2-2* characterizes a specific population of midbrain GABAergic neuron precursors

We next characterized the *Skor2*^+^ and *Nkx2-2*^+^ precursors in more detail. ISH analysis of parallel sections at E12.5 revealed that *Skor2* was expressed in a region coinciding with abundant GABAergic neuron precursors (Fig. 2A,B). Furthermore, the *Skor2^+^* post-mitotic precursors in the m2 co-expressed *Gad1* (Fig. 2D), and thus represent a subgroup of GABAergic neurons. Combined ISH and IHC analyses indicated that the *Skor2^+^* post-mitotic neuronal precursors in the m2 co-expressed *Nkx2-2* at E12.5 (Fig. 2E) and at E18.5 (Fig. 2G-H, J-K, M-M”). The *Skor2*, *Nkx2-2* and *Gad1* co-expressing cell population was found also in the WT adult midbrain (Fig. 2L). Similar to E12.5, *Skor2* expression was not detected in the *Gata2^cko^* midbrain at E18.5 (Fig. 2B-C, H-I), arguing for a fate change and against a delayed differentiation of these neurons in the absence of *Gata2* function. Thus, the co-expression of *Skor2* and *Nkx2-2* marks a highly restricted subgroup of Gata2 dependent midbrain GABAergic neurons.

To study the kinetics of differentiation of the *Skor2*^+^ *Nkx2-2*^+^ GABAergic neuron population, we determined the timing of terminal mitosis of the progenitors of these cells. For this, thymidine analogs EdU and BrdU were given at two consecutive days, between E9.5 and E12.5 and their incorporation was analyzed at E13.5. Thymidine analog injection at E9.5-E11.5 efficiently labelled the *Skor2*^+^ *Nkx2-2*^+^ cells, while injection at E12.5 produced only few scattered labelled cells in the posterior midbrain (Supplementary Fig. S2). These data suggest that *Skor2*^+^ *Nkx2-2*^+^ GABAergic neurons are mostly born between E11.5 and E12.5 and that their cell cycle exit may proceed in an anterior to posterior sequence. The birth-dating results are consistent with our gene expression studies that first detected both *Skor2* and *Nkx2-2* expression in the m2 region around E12.0, soon after the terminal mitosis occurs. These results also suggest that the *Skor2*^+^ *Nkx2-2*^+^ GABAergic precursors do not undergo tangential migration, but likely differentiate from the adjacent neuroepithelial progenitors.

### *Nkx2-2* is required upstream of *Skor2* for the subtype specification of midbrain GABAergic neurons

Next, we asked if *Nkx2-2* and *Skor2* are required for the differentiation of midbrain GABAergic neurons. We first analyzed *Nkx2-2* null mutant mouse embryos homozygous for a *Nkx2-2^Cre^* allele (*Nkx2-2^Cre/Cre^*), combined with an *Ai14^TdTomato^* reporter allele, which allowed us to follow the development of the mutant cells. At E12.5, we detected a reporter-labelled cell cluster in the m2 domain, both in the *Ctrl* (*Nkx2-2^Cre/+^*; *Ai14^TdTomato/+^*) and *Nkx2-2^null^* (*Nkx2-2^Cre/Cre^*; *Ai14^TdTomato/+^*) embryos (Fig. 3A-C, A’-C’). While in the *Ctrl* midbrain the labelled precursors expressed *Skor2*, we could not detect *Skor2* expression in the labelled precursors in the *Nkx2-2^null^* (Fig. 3 D-D’). In contrast to the midbrain, *Skor2* expression in the rhombomere 1 of the *Nkx2-2^null^* embryos was not affected (data not shown). Similar to E12.5, *Skor2* expressing cells were not detected in the midbrain of *Nkx2-2^null^* embryos at E18.5 (Fig. 3I-I’, J). This argues against delayed activation of *Skor2* in the *Nkx2-2* deficient precursors, and suggests that Nkx2-2 is functionally required for *Skor2* transcription in the m2.

**Figure 3.**
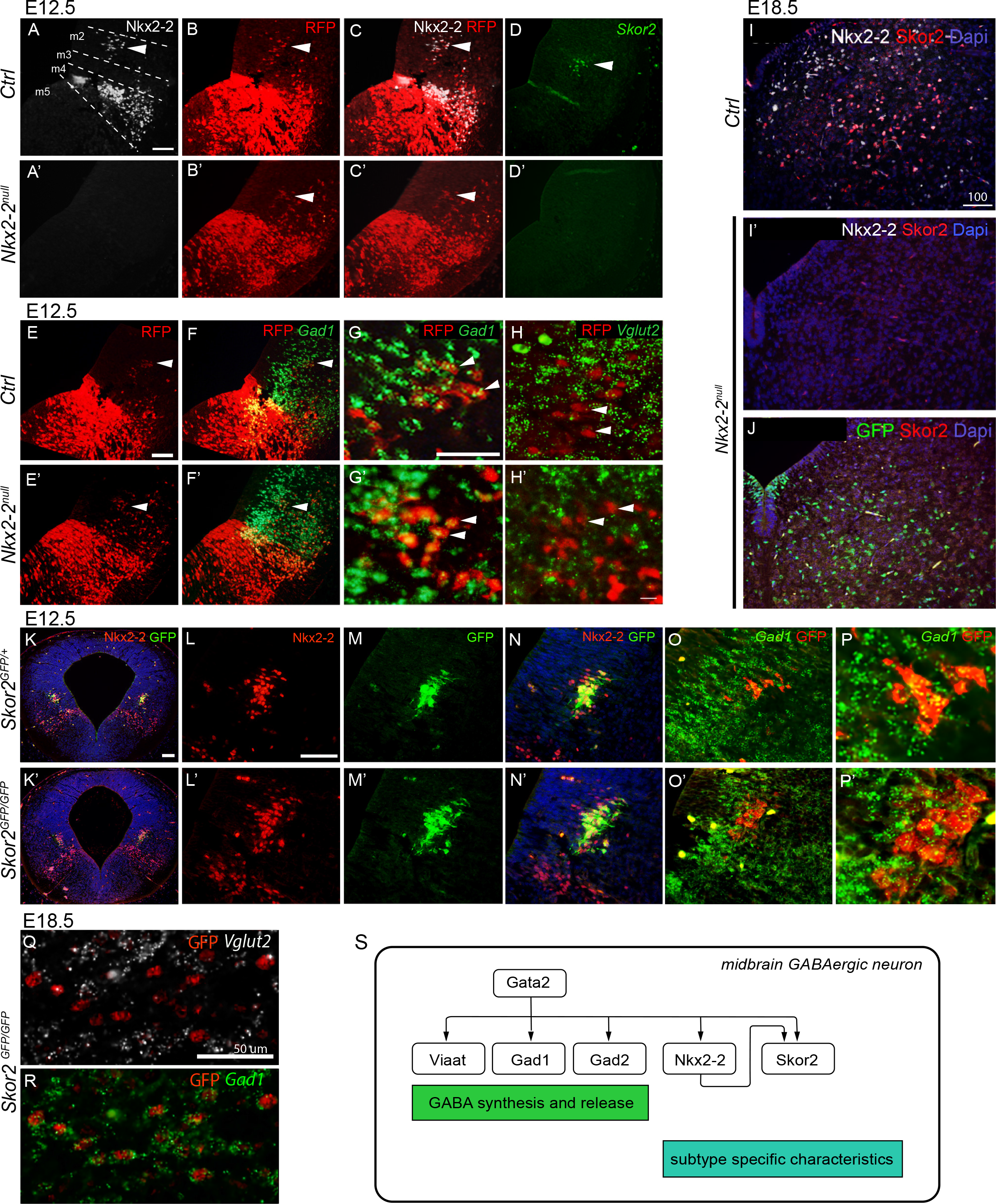
The function of *Nkx2-2* and *Skor2* in the developing midbrain GABAergic precursors. A-D’, Nkx2-2 is required for *Skor2* expression in the m2. IHC for Nkx2-2 and RFP in the E12.5 *Ctrl* (*Nkx2-2-^cre/+^, Ai14^TdTomato /+^*) and *Nkx2-2^null^* (*Nkx2-2^cre/cre^, Ai14^TdTomato /+^*) midbrain (A-C’) indicates that Nkx2-2 lineage precursors, marked by RFP expression, are generated in the midbrain m2 domain of the *Nkx2-2^null^* embryos, but they lack *Skor2* expression (ISH) (arrowheads). E-H’, Expression of *Gad1* (F-G’) and *Vglut2* (H,H’) in the *Nkx2-2* lineage cells is unaltered by the loss of Nkx2-2 function. ISH for *Gad1* or *Vglut2* combined with IHC for RFP in the *Ctrl* (*Nkx2-2-^cre/+^, Ai14^TdTomato /+^*) and *Nkx2-2^null^* (*Nkx2-2^cre/cre^, Ai14^TdTomato /+^*) midbrain at E12.5. I-I’, Expression of Skor2 (IHC) and Nkx2-2 (IHC) in E18.5 *Ctrl* and *Nkx2-2^null^* midbrain. Skor2 expression is lost in the *Nkx2-2^null^* midbrain. J, Expression of Skor2 (IHC) and *Nkx2-2* (IHC for GFP transcribed from the *Nkx2-2^Cre^* allele, where both Cre and EGFP sequences are inserted into the *Nkx2-2* locus) in E18.5 *Nkx2-2^cre/cre^* midbrain. Cells expressing the *Nkx2-2* gene are still detected but they are negative for Skor2. K-N’, Co-expression of Nkx2-2 (IHC) and *Skor2* (IHC for GFP encoded by the *Skor2^GFP^* allele) in E12.5 *Skor2^GFP/+^* and *Skor2^GFP/GFP^* midbrain. O-P’, Co-expression of *Gad1* (ISH) and *Skor2* (IHC for GFP encoded by the *Skor2^GFP^* allele) in E12.5 *Skor2^GFP/+^* and *Skor2^GFP/GFP^* midbrain. Q-R, Analysis of GFP (IHC), *Vglut2* (ISH) and *Gad1* (ISH) expression in the midbrain of E18.5 *Skor2^GFP/GFP^* embryo, demonstrating the maintenance of a GABAergic identity by the *Skor2* mutant cells. S, A schematic model of the transcriptional regulatory network driving differentiation of the *Skor2* expressing GABAergic neuron subtype. Scale bars: 200 μm (K, K’), others 50 μm.

To study the neurotransmitter identity of the m2 precursors in the absence of Nkx2-2, we analyzed the midbrain of E12.5 *Nkx2-2^null^* embryos for the expression of the GABAergic neuron markers Gata3 and *Gad1* and glutamatergic neuron marker *Vglut2 (Slc17a6*). The m2 precursors labelled by *Nkx2-2^Cre^* expressed *Gad1*, but not *Vglut2*, both in the *Ctrl* and the *Nkx2-2^null^* embryos (Fig. 3E-H, E’-H’). Thus, in the absence of Nkx2-2 function, the m2 precursors still acquire a GABAergic identity. However, the GABAergic subtype identity of m2 derivatives may still be altered as the *Skor2* expression is lost.

To study if Skor2 is required for the *Nkx2-2* expression and differentiation of the m2 precursors, we used mice carrying a *Skor2^GFP^* allele (Nakatani et al., 2014), where the coding sequences of *Skor2* are replaced with EGFP. Neither *Skor2* mRNA, nor Skor2 protein was detected in *Skor2^GFP/GFP^* embryos, confirming the loss of Skor2 function. At E12.5, EGFP positive cells expressing Nkx2-2 and *Gad1* were found in the m2 domain in both *Skor2^GFP/+^* and *Skor2^GFP/GFP^* embryos (Fig. 3K-P, K’-P’) as well as E18.5. Thus, the prospective *Skor2/Nkx2-2* expressing cell lineage was maintained in the absence of *Skor2* function. Furthermore, the unaltered *Gad1* and *Vglut2* expression in E18.5 shows that also the GABAergic identity was maintained in the *Skor2^GFP/GFP^* cells (Fig. 3Q,R).

Together, these results show that in the differentiating midbrain m2 precursors, Nkx2-2 is required upstream of Skor2. The cell cycle exit, cell survival, and GABAergic neurotransmitter fate specification of the m2 precursors appear independent of Nkx2-2 and Skor2. Instead, these TFs might regulate acquisition of GABAergic subtype-specific neuronal characteristics (Fig. 3S).

### *Skor2* expressing precursors give rise to GABAergic neurons at the boundary of the dMRF and vlPAG

The expression of *Skor2* and Nkx2-2 in a highly restricted population of embryonic midbrain GABAergic precursors likely signifies the development of an anatomically and functionally distinct subtype of GABAergic neurons. Therefore, we asked whether *Skor2*;*Nkx2-2* positive neurons are located in unique midbrain GABAergic nuclei in the postnatal brain.

We detected EGFP expression in the P4 and adult *Skor2^GFP/+^* mice, demonstrating specific *Skor2* expression in a subset of GABAergic neurons in the dMRF and in the adjacent vlPAG region of the midbrain (Fig. 4A-G). *Skor2*+ cells were found throughout the midbrain, with their amount decreasing caudally (Fig. 4B-D, H). Analysis of neurofilament and tyrosine hydroxylase expression further indicated localization of the *Skor2*+ cells at the dMRF/vlPAG boundary (Fig. 4E, F). In addition to the midbrain, *Skor2* was expressed in the cerebellar Purkinje cells and uncharacterized nuclei in the ventral hindbrain (Fig. 4H, Supplementary Fig. S4).

**Figure 4.**
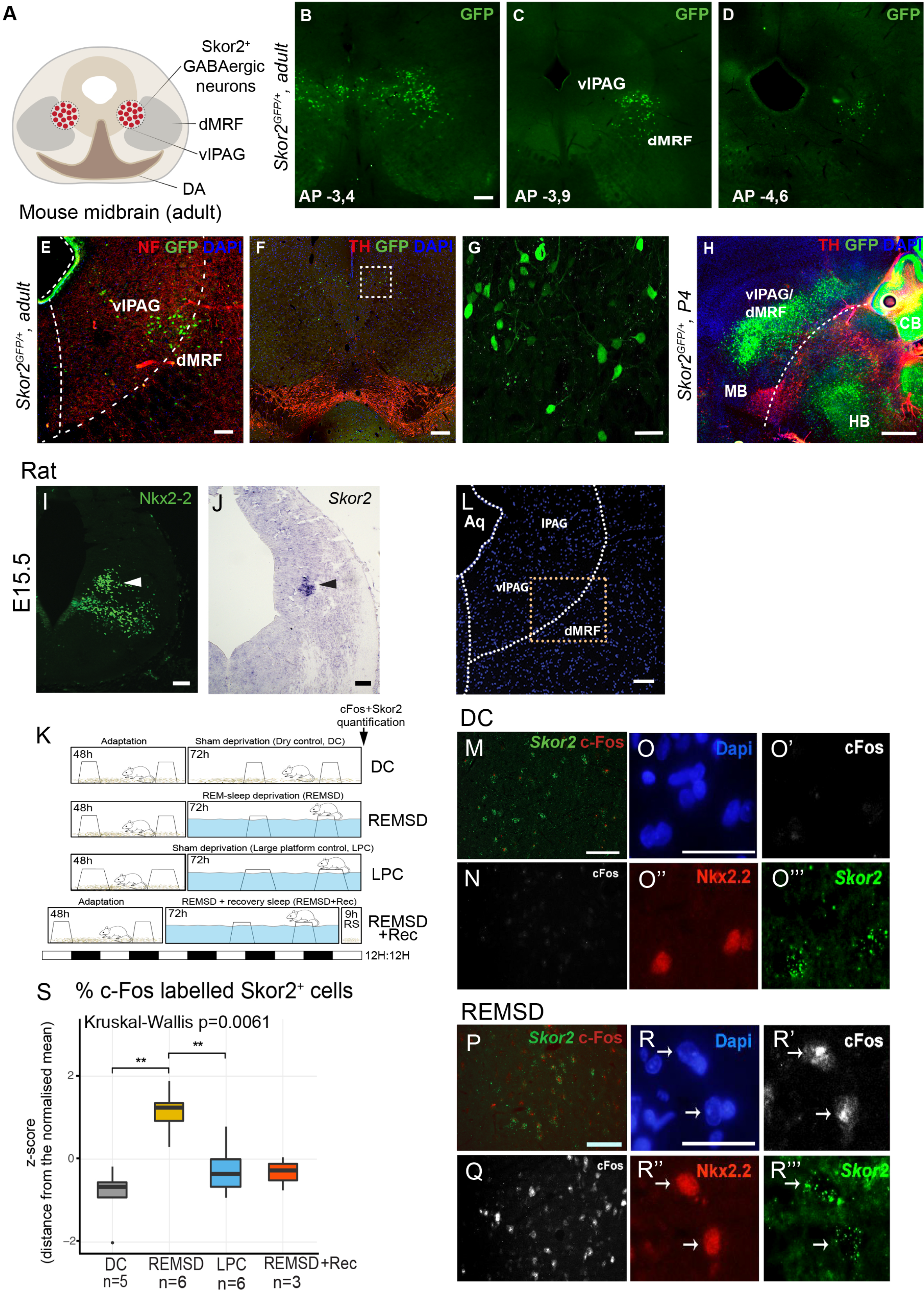
*Skor2* expressing neurons are located at the boundary of dMRF and vlPAG in the adult brain and their activity is increased by REM sleep deprivation. A, Schematic coronal view to the mouse midbrain, indicating the position of the *Skor2*^+^ cells. B-D, GFP expression in coronal sections of the adult *Skor2^GFP/+^* mouse brain. The distance from the bregma is indicated on each section. Scale bar, 100 μm E-G, IHC analysis of GFP expression in relation to neurofilament (NF, E) and TH (F) expression in the adult *Skor2^GFP/+^* mouse midbrain. Coronal section of the midbrain vlPAG and dMRF regions. G, Close-up of the *Skor2*^+^ neurons located in the area indicated with the dashed square in F. Scale bars, 100 μm, 50μm H, Sagittal section of the P4 *Skor2^GFP/+^* mouse midbrain analyzed for the expression of TH and GFP (IHC). Scale bar, 500μm I-J, Nkx2-2 (IHC) and *Skor2* expression (ISH) on coronal sections of the E15.5 rat midbrain. K, Schematic presentation of the REM-sleep deprivation assay. The experiment groups, treatment times and conditions are indicated. L, A coronal section of the adult rat midbrain showing the area analysed for the *Skor2* and c-Fos expression (vlPAG/dMRF). M-R’’’, Images of the c-Fos (IHC), Nkx2-2 (IHC) and *Skor2* (ISH) expression in the dry control (DC) and REM-sleep deprived rats (REMSD). All sections are from the dMRF area; low-magnification and a close-up is shown per area and experimental group. c-Fos+, Skor2+ cells were counted from the co-stained sections covering the whole dMRF area, to obtain data for the quantifications shown in S. S, The proportion (%) of *Skor2*^+^ cells that express c-Fos in the REM-sleep deprived rats (REMSD) and the control groups: rats in dry tank with platforms (DC), large platforms (LPC), and rats allowed a recovery period for 9 hours after REM sleep deprivation (REMSD+Recovery). Scale bars, 50 μm.

### REM sleep deprivation activates the *Skor2* expressing neurons in the dMRF/vlPAG

GABAergic neurons at the boundary of the dMRF and the vlPAG have been implicated in inhibition of the REM sleep and control of transitions between REM and non-REM sleep (Boissard et al., 2003; Hayashi et al., 2015; Lu et al., 2006; Weber et al., 2018). The activity of these REM-off neurons is increased by experimental REM sleep deprivation, as demonstrated by upregulation of c-Fos expression (Sapin et al., 2009). As the anatomical location of the REM-off neurons appears very similar to the location of *Skor2* and *Nkx2-2* expressing neurons at the dMRF/vlPAG boundary, we hypothesized that the *Skor2*^+^ *Nkx2-2*^+^ neurons represent the REM-off neurons. To test this, we asked if c-Fos expression was affected in the *Skor2* expressing cells by REM sleep deprivation. For this, we implemented the REM-sleep deprivation model using the inverted flowerpot method established for the rat (Sapin et al., 2009) (Fig. 4K). We first verified that, similar to the mouse, the *Skor2* and *Nkx2-2* expression is specific to the embryonic m2 mantle zone and adult dMRF/vlPAG in the rat (Fig. 4 I-J, L, M, P). We then analyzed the co-expression of c-Fos and *Skor2* in the dMRF/vlPAG in the REM sleep deprived rats compared to the control groups or recovery group (Fig. 4L-R’’’). The comparison of the the proportion of dMRF/vlPAG *Skor2*^+^ neurons expressing c-Fos revealed a significant increase in the c-Fos expression in the REM-sleep deprived rats (n=6, 72 hours deprivation of REM sleep) compared to the control groups (DC, n=6; LPC, n=5), and a recovery group (n=3, 72 hours REM-sleep deprivation followed by 9 hours period of normal sleep conditions) (Fig. 4S, Kruskal-Wallis H=12.399, p=0.0061; Supplementary Fig. S5 and Supplementary Table S5).

**Figure 5.**
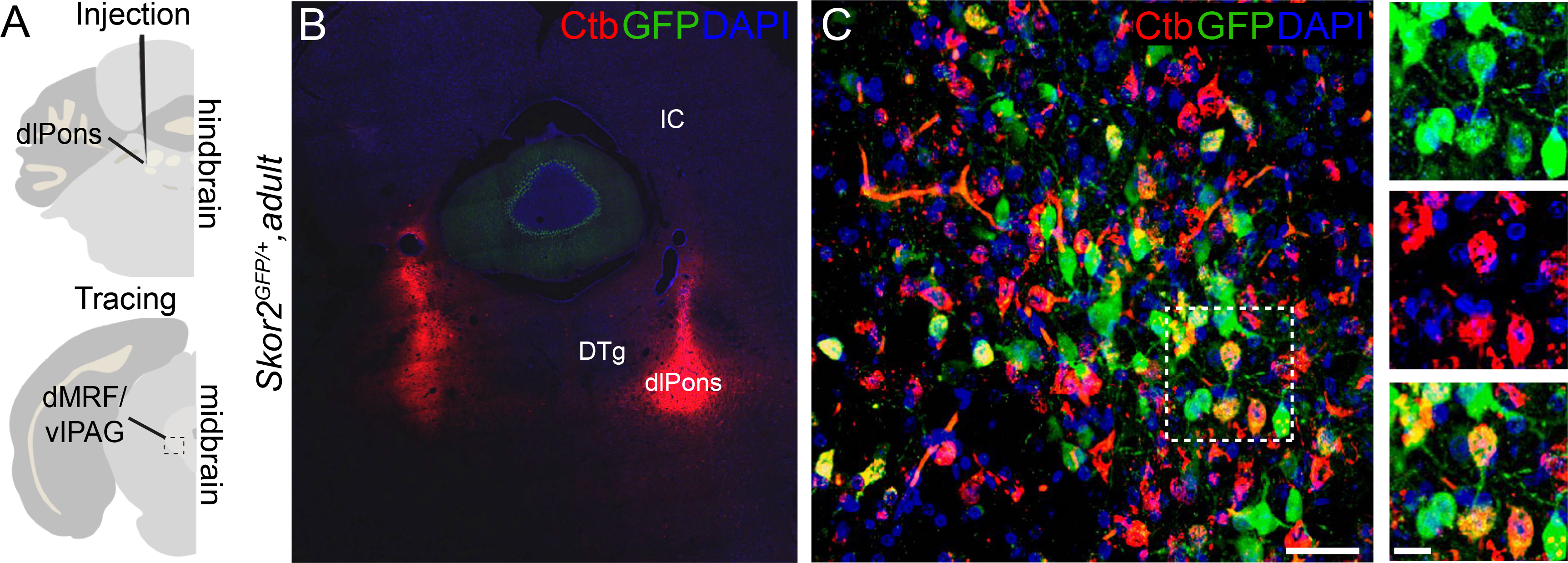
Retrograde labeling of the dMRF/vlPAG *Skor2*^+^ neurons from the dorsolateral pons. A, Experiment design for the retrograde labelling. CtB tracer was injected in the dorsolateral pons (dlPons) in hindbrain (n=6). dMRF/vlPAG area in the midbrain was analyzed for the tracer expression. B, Injection site after CtB injection. DTg, dorsal tegmental nucleus. Scale bar, 50 μm. C, dMRF region in the injected animal analysed for the co-expression of GFP and CtB (IHC). Examples of double labelled cells are shown in the close-up in the right (white box). Scale bar, 10 μm.

### *Skor2* expressing neurons in the dMRF/vlPAG project to the dorsolateral pons

Earlier studies have shown that dMRF/vlPAG GABAergic neurons project to the dorsolateral pons and inhibit its REM sleep promoting activity (Boissard et al., 2003; Hayashi et al., 2015; Lu et al., 2006; Sapin et al., 2009; Weber et al., 2018). To test whether the *Skor2* expressing GABAergic cells in the dMRF/vlPAG project to the dorsolateral pons, we used retrograde labelling. Retrograde tracer Choleratoxin subunit B (CtB) was injected to the dorsolateral pons of the adult *Skor2^GFP/+^* mice (Fig. 5A, B). In all the injected animals (n=6), CtB labelled cells in the dMRF/vlPAG region of the midbrain (Fig. 5C), consistent with dorsolateral pons receiving inputs from the dMRF. Many of the CtB labelled cells showed co-expression of *Skor2* (Fig. 5C, close-ups), suggesting that the *Skor2^+^* dMRF/vlPAG neurons frequently project to the dorsolateral pons.

### *Skor2^+^* neurons express functional Orexin receptors

Orexinergic signalling via the Orexin receptors (Hcrtr1 and Hcrtr2) regulates sleep and wakefulness, and loss of the hypothalamic orexinergic neurons is associated with narcolepsy with cataplexy, possibly due to altered input to vlPAG GABAergic neurons that express Orexin receptors (Kaur et al., 2009; Lu et al., 2006). We analysed the expression of Hcrtr1 and Hcrtr2 in *Skor2* expressing cells in the dMRF/vlPAG of the adult *Skor2^GFP/+^* mice (n=4) by IHC (Fig. 6A-B). We found that of the *Skor2*^+^ cells expressing GFP, 68.5% (sd=1.6) expressed Hcrtr1 and 69.5% (sd=0.7) expressed Hcrtr2. Since the proportion of both *Skor2*^+^/Hcrtr1^+^ and *Skor2*^+^/Hcrtr2^+^ double positive neurons were above 50% in all animals (n=4), the co-expression of both receptors is likely, but not analysed here.

**Figure 6.**
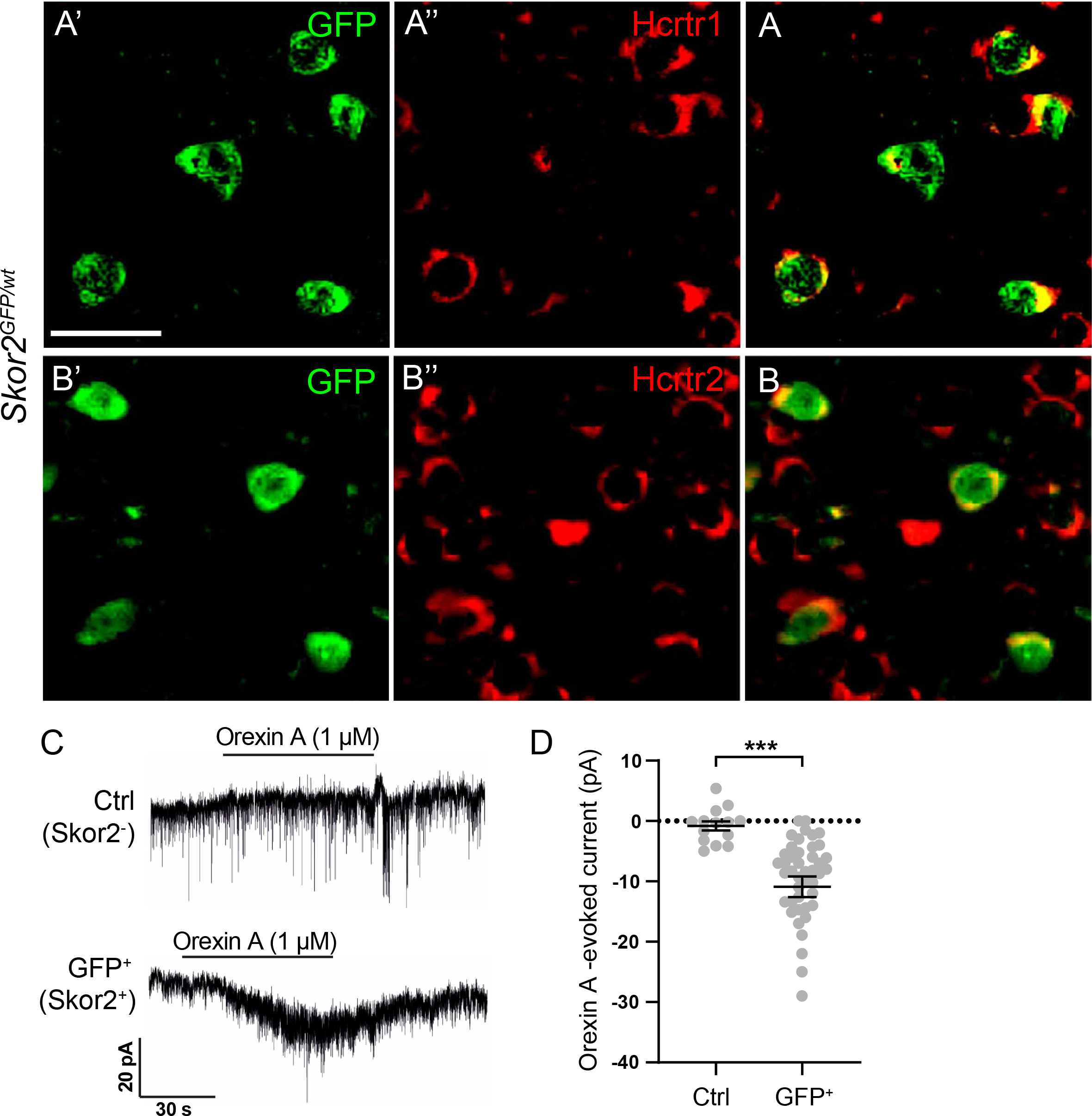
Orexin receptor expression and Orexin-evoked currents in the *Skor2*^+^ neurons. A-B, IHC analysis of the co-expression of GFP and Hcrtr1 (A) or Hcrtr2 (B) in the dMRF/vIPAG region of the adult *Skor2^GFP/+^* mice. Scale bars, 50 μm. C, Representative traces of holding currents during Orexin A application in GFP-negative (control) and GFP-positive (*Skor2***^+^**) cells. D, Summary of Orexin A induced currents (individual values and mean±SEM) in GFP-negative (control, n=14 cells, 3 animals) and GFP-positive (*Skor2***^+^**, n=44 cells, 9 animals) cells. *** P<0.001 (one-way ANOVA on ranks).

We next asked whether the *Skor2*^+^ cells respond to Orexin. In voltage clamp recordings of acute midbrain slices of adult *Skor2^GFP/+^* mice, bath application of Orexin A (1 µM) induced an inward current in the GFP expressing *Skor2^+^* cells, but not the neighbouring control cells (-10.9 ± 1.7 pA vs -2.2 ± 0.9 pA, p<0.001, Fig. 6C-D). The proportion of responding cells (Skor2^+^: 32/44 cells, 72 %; Skor2^-^: 0/14 cells; Fig. 6D) correlated with the proportion of the *Skor2^+^* cells expressing Orexin receptors.

### Functionally and anatomically distinct subtypes of *Skor2*^+^ neurons

Finally, we characterized the firing properties of the *Skor2*^+^ cells in the dMRF/vlPAG region in acute midbrain slices of adult *Skor2^GFP/+^* mice (n=17) by whole cell patch clamp. We analysed passive and active membrane properties of 42 GFP expressing *Skor2*^+^ cells, and saw a clear difference in patterns of action potential firing. Because of the heterogeneity of the neurons, we decided to divide the cells into three subgroups, referred to here as *adapting* (n=13), *stuttering* (n=14) and an *intermediate* neurons (n=15) (Fig. 7, Supplementary Fig. S5). Cells were assigned to “stuttering” group, if they displayed fast-decaying afterhyperpolarizing potential (AHP) and exhibited characteristic stuttering action potential firing with bursts of action potentials (APs) intermingled with quiescent periods. Quiescent period was defined as an interval, which is at least three times longer than the preceding and consequent intervals. The rest of cells had AHP with longer decay and fired action potentials in “adapting” mode, with regular AP intervals becoming longer towards the end of the current step. Cells were assigned to “adapting” group, if their AHP consisted of two clear components, separating fast and medium AHP. The rest of the cells, displaying adapting firing and AHP without two components, was placed to the “intermediate” group. The division of cells into three subgroups was supported by statistically significant difference in individual parameters of AP firing. Adapting neurons had the highest rate of spontaneous activity (adapting: 7.5 ± 1.4 Hz, stuttering: 0.7 ± 0.3 Hz, intermediate: 4.7 ± 1 Hz) and the lowest rheobase (adapting: 31.5 ± 6 p*A,* stuttering: 92.1 ± 13.7 pA, intermediate: 56.7 ± 7.9 pA) (Fig. 7E-H). The neuron classes also significantly differ in the amplitude of the medium AHP (adapting: 15.1 ± 0.8 mV, stuttering: 7.4 ± 0.7 mV, intermediate 12.3 ± 0.9 mV) and voltage response to hyperpolarizing current steps (slope of the linear IV relationship, adapting: 0.6 ± 0.1 mV/pA, stuttering: 0.3 ± 0.04 mV/pA, intermediate: 0.4 ± 0.05 mV/pA). Other passive and active membrane properties were similar (Supplementary Fig. S5). The kinetics of the afterhyperpolarization, as well as hyperpolarization-induced currents suggest a differential expression of unknown K^+^ channels in the *Skor2*^+^ neuron subgroups.

**Figure 7.**
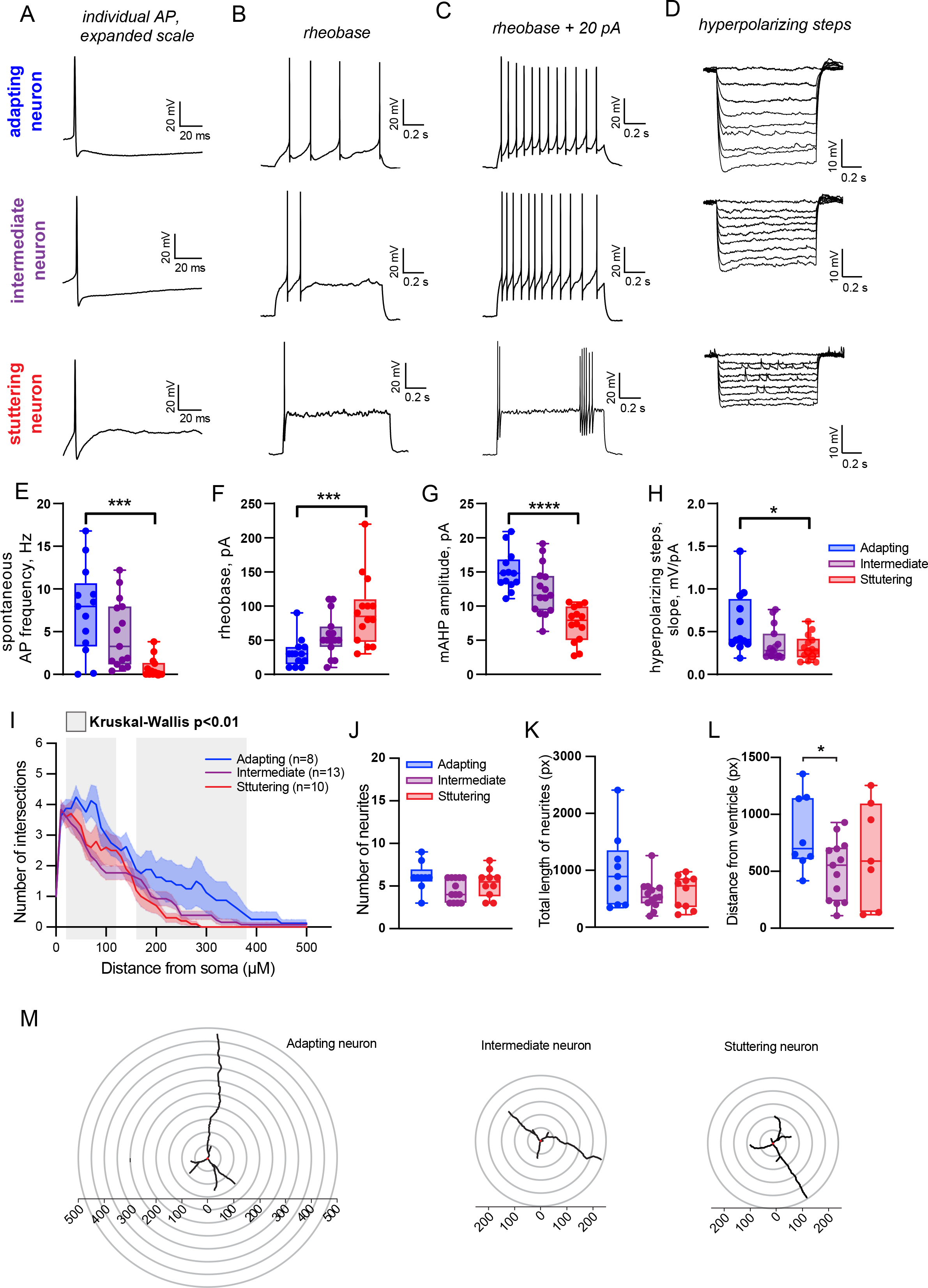
Three subclasses of dMRF/vlPAG *Skor2*^+^ neurons with distinct electrophysiological and morphological characteristics. A-D, Example traces of evoked action potential firing and voltage response to hyperpolarization in different types of *Skor2*^+^ cells in the mouse midbrain. E-H, Excitability parameters of the *Skor2*^+^ cells. Data are presented as individual values and box plot showing median, 25 and 75 percentiles with whiskers showing minimum and maximum values. AP: action potential; mAHP: medium after-hyperpolarizing potential. Adapting cells n=13; intermediate cells n=15, and stuttering cells n=14, 17 animals. *p<0.05; ***p<0.001; ****p<0.0001, one-way ANOVA. I, Sholl analysis comparing intermediate (n= 8), adapting (n= 13) and stuttering (n= 10) *Skor2*+ cells for the number of intersections at a various distance from soma or cell body (16 animals). Data represented as mean±SEM. The gray shading indicates the region where the three groups show statistically significant differences (Kruskal-Wallis p<0.01). J-K, Quantification of the neurite number (J) and total length of neurites (K) in adapting, intermediate and stuttering type of *Skor2*^+^ neurons. Neurite number and total (summed) neurite length was calculated from the Sholl traced neurons. L, Comparison of the distance of the cell soma from the midbrain ventricle in the three subclasses of *Skor2*+ neurons. A difference between the means of distance from the ventricle was detected between adapting and intermediate neuron types (t-test p<0.05). M, Reconstructions or tracing images of representative neurons used for Sholl analysis.

To morphologically characterise the *Skor2*^+^ cells, we filled the recorded cells with biocytin, allowing visualization of their neurites in the midbrain slices. Sholl analysis of the biocytin-filled cells (n=44, cells from 16 animals) suggested that, as a group, the GFP expressing *Skor2*^+^ neurons (n=35) did not significantly differ from the neighbouring neurons negative for GFP expression (n=9, Supplementary Fig. S7A, B-C). However, we detected differences among the *Skor2*^+^ neuron subclasses with different firing patterns. The adapting neurons exhibited slightly higher number of intersections compared to intermediate or stuttering neurons (Fig. 7I). The most significant differences were observed between adapting and intermediate neurons in a zone 20–120 µm and 170-380 µm from the soma (Kruskal-Wallis p<0.01, Fig. 7I, Supplementary Fig. S7D-G). However, no statistically significant differences were detected in the total neurite numbers or total neurite lengths of the three *Skor2*^+^ neuron subgroups (Fig. 7J-K). The thickness of the slice (250 µm) likely limits these analyses. We also mapped the localization of the *Skor2*^+^ neurons relative to the ventricle. Our results suggest that, compared to the intermediate neurons, the adapting neurons are localised farther away from the aqueduct (Fig. 7L). Thus, the adapting neurons show the longest projections and appear to localize primarily in the dMRF rather than in vlPAG.

In summary, the *Skor2*^+^ cells in the mouse dMRF/vlPAG region are morphologically and physiologically diverse. A large proportion of the *Skor2*^+^ cells are responsive to Orexin via Hcrtr1/2 and could thus represent the orexin-responsive neurons regulating sleep.

## Discussion

The central regions of the midbrain, the PAG and MRF, contain neurons important for regulation of multiple aspects of behavior, including defensive behaviors, motivated behaviors, attention and sleep. However, studies of these neurons are hampered by the lack of knowledge on their subtype-specific molecular features and developmental regulation. Gata2 acts as a selector gene required for the differentiation of all midbrain-derived GABAergic lineages, but the mechanisms of subtype specific fate regulation in those lineages are not resolved. We have here identified several Gata2-regulated and GABAergic subtype specific TFs, including Nkx2-2 and Skor2. We show that Gata2, Nkx2-2 and Skor2 mark and regulate the development of an anatomically restricted dMRF/vlPAG GABAergic neuron population potentially involved in regulation of the REM sleep.

### Gene regulatory hierarchy between Gata2, Nkx2-2 and Skor2

We showed that the differentiation of *Skor2* expressing GABAergic neurons is dependent on both *Gata2* and *Nkx2-2* gene function. In the ventral midbrain, including the m4 region, *Nkx2-2* is expressed in the ventricular zone progenitors as a part of the homeodomain TF code of patterning the ventral neural tube (Kala et al., 2009; Nakatani et al., 2007; Prakash et al., 2009; Puelles et al., 2004). In post-mitotic precursors derived from the m4, *Nkx2-2* expression is maintained after the cell cycle exit and expressed during the subsequent development of both GABAergic and glutamatergic neurons, which remain to be anatomically and functionally characterized. In contrast, in the m2, *Nkx2-2* and *Skor2* were activated only in the post-mitotic precursors in a Gata2 dependent fashion. With the genetic loss-of-function analyses, we demonstrated that the *Nkx2-2* null mutant embryos lose *Skor2* expression, while *Nkx2-2* expression is retained in the absence of *Skor2* gene function. Those results indicate the regulatory cascade where both *Nkx2-2* and *Skor2* genes are activated after the cell-cycle exit, when Gata2 directly or indirectly activates *Nkx2-2*, which in turn regulates the expression of *Skor2*. However, to demonstrate direct regulatory cascade, chromatin association experiments would be needed to test the binding of Gata2 on the *Nkx2-2* and Nkx2-2 on *Skor2* gene regulatory elements. Furthermore, the target genes of Skor2 TF in the vlPAG/dMRF lineage are currently unknown.

The generic GABAergic features of the midbrain precursors (such as the expression of *Gad1* and *Gata3*) seem to be unaffected in the absence of Nkx2-2 or Skor2 function. This is in contrast to cerebellar Purkinje cells, where Skor2 is required for the proper acquisition and maintenance of a GABAergic phenotype (Nakatani et al., 2014; Wang et al., 2011). In midbrain, the pan-GABAergic features of the m2 precursors are likely controlled by the Gata/Tal TF complex (Achim et al., 2013; Kala et al., 2009). Nkx2-2 and Skor2 may regulate further neuronal subtype-specific characteristics, such as neurotransmitter reception, excitability, connectivity patterns, cellular morphology, or co-neurotransmitter expression. Single-cell or cell type specific RNA-sequencing could provide more information on the molecular composition of the *Nkx2-2*^+^/*Skor2*^+^ GABAergic neurons.

### Skor2 as a marker of dMRF/vlPAG REM-off neurons

Interactions between REM sleep promoting REM-on neurons in the dorsolateral pons and REM sleep inhibiting REM-off neurons in the vlPAG/dMRF are thought to regulate normal sleep cycles (Luppi et al., 2017; Saper et al., 2010; Scammell et al., 2017; Weber and Dan, 2016). In rats, REM sleep deprivation results in stimulation of dMRF/vlPAG GABAergic neurons, as evidenced by upregulation of the expression of the immediate-early gene c-Fos (Sapin et al., 2009). A recent study of neuronal activity during sleep episodes demonstrated that dMRF/vlPAG GABAergic neurons increase their firing during transitions from non-REM sleep to wakefulness also in mice, consistent with the suggested REM-off activity (Weber et al., 2018). Optogenetic and chemogenetic activation of the dMRF/vlPAG GABAergic neurons decreases REM sleep, whereas their inhibition increases

REM sleep (Hayashi et al., 2015; Weber et al., 2018). An important projection target of the inhibitory dMRF/vlPAG REM-off neurons is the dorsolateral pons, which contains excitatory glutamatergic REM-on neurons controlling both muscle atonia and other aspects of REM sleep. In turn, the regulatory inputs into dMRF/vlPAG region include orexinergic excitatory projections from the ventral hypothalamus, which are implicated in the narcolepsy and narcolepsy-associated loss of muscle tone reminiscent to REM sleep (Kaur et al., 2009; Lu et al., 2006; Willie et al., 2003). The *Skor2*^+^ neuron population described here seems to be integrated in this circuit in both directions.

Although their role in REM sleep regulation is well established, the GABAergic neurons in the dMRF/vlPAG remain a heterogeneous population, containing cells of both REM-off and REM-on activity (Luppi et al., 2017; Weber et al., 2018; Verret et al., 2006). We show here that the location of the *Skor2* expressing dMRF/vlPAG cells matches the position of the REM-off neurons in the mouse and rat. Upregulation of c-Fos upon REM sleep deprivation, projection to dorsolateral pons, and responsiveness to orexin further support the hypothesis that at least some of the *Skor2* positive cells represent dMRF/vlPAG REM-off neurons. Differences in the electrophysiology profiles and neuronal morphology of the *Skor2* expressing neurons may indicate the presence of additional functional subclasses, which could have distinct roles in sleep regulation or other dMRF/vlPAG mediated functions, such as non-REM sleep stages, eye and body movements, and nociception. Recording and modulation of the activity of the *Skor2* expressing neurons during sleep behavior will be needed to rigorously test their involvement in REM sleep regulation. As the homozygous *Skor2* mutant mice die shortly after birth (Nakatani et al., 2014), conditional inactivation of *Skor2* in the midbrain will be needed to test the requirement of this TF for distinct subtype-specific anatomical and physiological properties of the dMRF/vlPAG GABAergic neurons.

### Other cell types and functions of the MRF and PAG

In addition to the m2 derived *Skor2* expressing GABAergic neurons described here, the MRF and PAG contain variety of other cell types. Some of these are also involved in sleep regulation. For example, the vlPAG contains a group of Neurotensin-expressing glutamatergic neurons that control non-REM sleep (Zhong et al., 2019). Furthermore, in addition to the GABAergic projection, the dMRF/vlPAG has been suggested to send glycinergic and glutamatergic projections to the dorsolateral pons including the sublaterodorsal nucleus (Liang et al., 2014). Studies of the derivatives of the other embryonic midbrain regions, including the Nkx2-2 expressing GABAergic and glutamatergic precursors in the m4, potentially give insights into the molecular, anatomical and functional diversity of these neurons.

Our study of the developmental and molecular properties of the dMRF/vlPAG neurons illustrates that unique molecular markers of GABAergic neuron subtypes can be identified, and that such markers facilitate studying the function of specific neuronal circuits. Single-cell transcriptomic analyses would help to further dissect the heterogeneity the developing midbrain region and the neuronal circuits therein.

## Materials and methods

### Mouse lines

*En1^Cre^* (Kimmel et al., 2000), *Gad67^GFP^* (Tamamaki et al., 2003), *Gata2^flox^* (Haugas et al., 2010), *Ai14^TdTomato^* (Madisen et al., 2010) were on an outbred (ICR) background and *Nkx2-2^Cre^* (Balderes et al., 2013) and *Skor2^GFP^* (Nakatani et al., 2014) alleles were maintained on a mixed background (C57BL/6 and ICR). E0.5 was defined as noon of the day of the vaginal plug. Experiments were approved by the Laboratory Animal Center, University of Helsinki, and the National Animal Experiment Board in Finland.

### Microarrays

Ventral and dorsal midbrain was dissected from E12.5 wild-type and *Gata2^cko^* (Kala et al., 2009) embryos. For both genotypes, 3 groups were generated, each consisting of 6 tissue samples. Total RNA was extracted with Trizol reagent and used for probe labelling. Illumina BeadChip (Mouse WG-6 2.0) microarrays were hybridized according to the manufacturer’s protocol. The data set was normalized using the quantile normalization method. Statistical testing was performed using LIMMA package using R and Bioconductor statistical analysis software. DAVID Bioinformatics Resources 6.7, NIAID/NIH (Huang da et al., 2009b; Huang da et al., 2009a) were used for the GO term enrichment analyses.

### Histology, mRNA in situ hybridization and immunohistochemistry

Dissected embryos, or brain tissue from embryos older than E16, were fixed in 4 % paraformaldehyde (PFA; Sigma-Aldrich P6148) in PBS. For adult mouse and rat brains intracardial perfusion was performed first with PBS and then with 4% PFA, and postfixed in 4% PFA. For paraffin embedding and microtome sectioning, samples were processed in Histosec polymer wax (Merck Millipore) and sectioned at 5 or 10 μm. For vibratome sectioning, adult mouse brain tissue samples were embedded in 4% agarose and sections were cut using vibratome (Leica VT1200S). Until further processing sections were stored in ice-cold 1xPBS. For in situ mRNA hybridization (ISH), digoxigenin (DIG) - labelled antisense cRNA probes were used. For ISH signal detection, tyramide signal amplification (TSA) -based method (TSA Plus Cy3 NEL744001KT /Fluorescein NEL741001KT; Perkin Elmer) was used for fluorescent detection and AP-based method for colorimetric detection.

For combined ISH and immunohistochemistry (IHC), ISH signal was visualized first followed by incubation with primary antibodies for IHC staining. For double ISH, DIG and fluorescein -labelled probes were combined. TSA Plus Cyanine 3 and Fluorescein kits were used for detection (protocols provided upon request). Antibodies and mRNA ISH probes are listed in Supplementary Table S4. For the rat Skor2 probe, rat Skor2 cDNA fragment (GeneArt sequence-based gene synthesis, Life Technologies) was cloned into pBluescript SK+ vector digested with NotI and SalI. The plasmid map and sequence are available upon request.

All analyses were confirmed using at least 3 biological replicates.

### Stereotaxic injections and retrograde tracing

Mice used in the experiments were S*kor2^GFP/wt^* (n=6). Mice were anesthetized with isoflurane, attached to the stereotaxic frame, and a small hole was drilled into their skull. For retrograde tracing of vlPAG neurons, bilateral, intracranial injections of 0, 2% Choleratoxin B subunit (#104; List Biological Lab.Inc.) were injected at the speed of 50 nl/min using a microinjector (UltraMicroPump III, World Precision Instruments) and microsyringe (Hamilton 7803-06). The stereotaxic coordinates for injections were (measured from bregma, in mm): -5,19 to -5,4 (AP); 0,88 (ML); -4,4 (DV). The coordinates were obtained from the mouse brain atlas (Paxinos and Franklin, 2012). Mice were intracardially perfused 5-7 days after the injections and the brains were collected. Brains were processed for vibratome sectioning and further for IHC stainings and analysis for GFP and CtB expression in the midbrain.

### REM sleep deprivation assay in rats

The REM sleep of male Han-Wistar rats was deprived for 72 h similarly as described previously (Porkka-Heiskanen et al., 1995) using the water tank (inverted flowerpot) method, when the animal has to sleep on a small platform surrounded by water. The platform is so small that the animal is not able to maintain its balance on it during the REM sleep-associated muscle hypotonia and falls into the water, thus REM sleep is suppressed almost totally.

Small platforms (inverted flowerpots, diameter: 6.5 cm) were placed into a round shape wire mesh cage situated in a basin. The wire mesh cage was provided with food tubes and water bottles. All animals were kept in in a 12h:12h light-dark cycle (light was on from 8:30 – 20:30) before and during the experiment. Before the REM sleep deprivation, the rats were placed into the dry apparatus (water tank) for 48 h in order to adapt them to the new environment. Then the basin was filled with water for 72 h. Simultaneously, 3 rats were placed into the wire mesh cage with 4 platforms. The animals had food and water *ad libidum*. The rats of the 2 control groups (Large platform control (LPC): large, 11-cm-diameter platforms were placed into the basin filled with water, the platforms were large enough to have REM sleep on them; Dry control (DC): there was no water in the basin with platforms, bedding was placed on the bottom) were kept in the apparatus for the corresponding time (48+72 h). An additional group of rats were allowed to have 9 h recovery sleep after the 72-h REM sleep deprivation (REMSD+Recovery group). The REM sleep deprivation of DC, LPC and REMSD rats started and ended 20-40 min before dark onset. The REM sleep deprivation of REMSD+Recovery group started end ended 9 h earlier (9 h 20-40 min before dark onset, i.e. 2 h 20-40 min after light onset). At the end of the REM sleep deprivation/sham deprivation or rebound sleep, the animals were sacrificed by intraperitoneal administration of 400 mg/kg chloral hydrate, perfused with PBS and then with 4% PFA, and the brains were removed for histochemical examination and for measuring gene expression.

For quantification of the c-Fos labelling, sections covering the dMRF/vlPAG area were collected from all REM-sleep assay animals and stained for the expression of c-Fos (IHC) and *Skor2* (ISH). The number of *Skor2*^+^ cells and the number of *Skor2*^+^ cells double labelled with c-Fos were registered (Supplementary Table S5).

Study contained two similar yet separate experiment series. These experiments series, while experiment design was identical, were conducted in different times. We observed systematic difference in c-Fos staining efficiency between experiments series (Supplementary Fig. S4). To alleviate this difference, we performed z-score transformation to get experiment series on the same scale. The scaled data was pooled before multiple comparison between groups. The number of animals in each group was: n=6 for REMSD, n=5 in the DC, n=6 in LPC, and n=3 in REMSD+Recovery group. The number of *Skor2*^+^ cells did not appear to vary between the experiments (*Skor2*^+^ cells; average N=72, sd=28). Kruskal-Wallis test for multiple comparison was used to test the variation in the mean transformed z-scores between all the experiment groups (experiment series merged). Pairwise group comparisons (experiment series merged) were performed using Wilcox-test.

The group identity, experiment ID, number of *Skor2*^+^ cells, number of c-Fos^+^ cells, the percent of c-Fos labelled *Skor2*^+^ cells and the transformed z-scores for each animal can be found in the Supplementary Table S5.

### Electrophysiology

Adult S*kor2^GFP/+^* mice of both genders (2-7 months old, n=27) were used for the preparation of acute brain slices. Animals were anesthetized with isoflurane. After decapitation, the brain was rapidly removed and transferred to ice cold cutting solution containing (in mM): 92 NMDG, 2.5 KCl, 1.25 NaH_2_PO_4_, 30 NaHCO_3_, 20 HEPES, 25 glucose, 2 thiourea, 5 Na-ascorbate, 3 Na-pyruvate, 0.5 CaCl2 and 10 MgSO_4_ (pH 7.3–7.4 was adjusted with concentrated hydrochloric acid). All extracellular solutions were equilibrated with 95% O_2_ and 5% CO_2_. Coronal slices (250 µm) containing the dMRF/vlPAG were cut using a vibrating microtome (7000 SMZ-2; Campden Instruments). Slices were kept for 10 min in NMDG cutting solution at 34 °C before being transferred to recovery solution at RT, containing (in mM): 92 NaCl, 2.5 KCl, 1.25 NaH_2_PO_4_, 30 NaHCO_3_, 20 HEPES, 25 glucose, thiourea, 5 Na-ascorbate, 3 Na-pyruvate, 2 CaCl2 and 2 MgSO_4_. Recordings were done 1-5 h after the preparation of acute slices. Whole cell patch clamp recordings from visually identified dMRF/vlPAG cells (GFP^+^ cells and neighboring GFP^-^ cells as control) were performed in a submerged recording chamber at 32 ± 0.5 °C, constantly perfused with ACSF (in mM: 124 NaCl, 3 KCl, 2 CaCl_2_, 26 NaHCO_3_, 1.25 NaH_2_PO_4_, 1 MgSO_4_, and 15 glucose, pH 7.4. Bath perfusion was 2.5 mL·min^−1^). Application of Orexin A (1µM, Tocris) and elevated K^+^ was done with a direct perfusion system locally onto the slice.

Whole-cell current-clamp recordings were obtained using a Multiclamp 700A patch-clamp amplifier and recorded with pClamp 10 (Molecular devices) at a sampling rate of 20–50 kHz. Borosilicate patch pipette resistance ranged from 3 to 6 MΩ. The composition of the patch pipette solution was the following (in mM): 135 K-gluconate, 10 HEPES, 5 EGTA, 4 Mg-ATP, 0.5 Na-GTP, 2 KCl, 2 Ca(OH)_2_, 280 mOsm (pH 7.2). The liquid junction potential of 13 mV was not corrected for. In some experiments biocytin (7 mM, Sigma) was included in the pipette solution to allow post hoc staining of the recorded cells.

For characterization of the cell properties, the cells were held in current clamp at -70 mV, and we injected hyperpolarizing and depolarizing current steps (1 s, increment of 10 pA). Excitability parameters were analyzed with the FFFPA script in Matlab (https://doi.org/10.5281/zenodo.3667731), except for the mAHP amplitude and hyperpolarizing steps. The mAHP amplitude was measured in Clampfit 11.1 as the mean voltage 15-20 ms after the AP threshold. The voltage response to current hyperpolarizing steps was measured in Clampfit 11.1. The hyperpolarizing steps slope was calculated by linear fit of voltage response amplitude plotted against the injected current. The effect of Orexin A on cells was assessed in voltage clamp. Cells were held at -70mV, and test pulses (-10 mV, 50 ms) were delivered every minute to monitor access resistance. The change in holding current was measured after 1 min of Orexin A application. For counting the number of cells which responded to Orexin application, the threshold for Orexin-evoked response was set at the level of baseline RMS noise multiplied by two. Placement of direct perfusion tubing was verified by application of 8 mM KCl at the end of experiment.

The analysis of electrophysiological recordings was done with Clampfit 11.1 (Molecular devices) and Matlab, statistical analysis was done with GraphPad Prism 9.

### Sholl analysis of neuronal morphology by Biocytin-filling

Biocytin filled GABAergic neurons from dMRF/vlPAG region were immunohistochemically stained and imaged using Leica SP8 STED confocal microscope. Tiled confocal z-stack images (40 images/stack) were acquired at 20x objective. Sholl analysis was performed as described (Comhair et al., 2018). Briefly, 8-bit images of dMRF/vlPAG GABAergic neurons were traced using the Simple Neurite Tracer (SNT v3.2.14) (Arshadi et al., 2021) plug in of FIJI (NIH, Bethesda, MD), and tracing files were generated. The number of dendrite and total dendrite length were generated using measurement function of Simple Neurite Tracer. The complexity of the neurites of the labelled neurons was evaluated using the Sholl analysis. To implement this, concentric sampling spheres with 10 μm intervals between the radii were formed around the central point, i.e., the soma of the traced neuron and a number of intersections with neurites was measured. For statistical analysis number of intersections per cell were used. Comparative analysis of Sholl curves between neuron types was made with Kruskal-Wallis test within 50 μm wide moving window. Distance intervals with statistically significant difference between the curves (p-value <0.01) are shaded in gray (Fig 7H). Separate pairwise comparison between Sholl curves of neuron types was done with Kolmogorov-Smirnov test within 50 μm wide moving window. Distance intervals with statistically significant difference between the curves were indicated (p-value <0.01). Distance from ventricle per biocytin-filled neuron types was measured and analysed by t-test. Normality of the distance distribution was tested with Shapiro before conducting the t-test. Statistical analyses were performed in GraphPad Prism 9 and R.

**Table 1.**
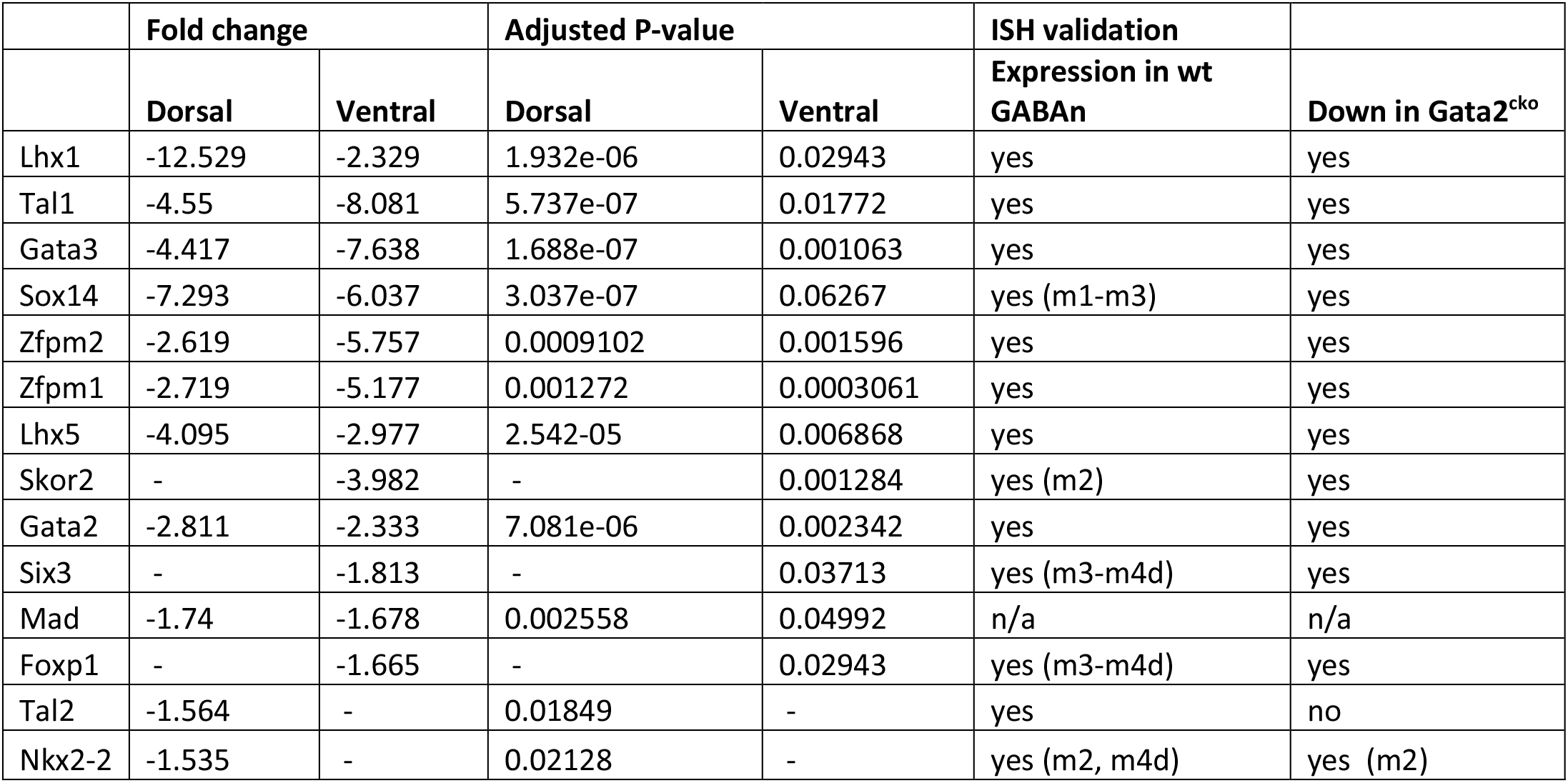
TFs down-regulated in Gata2 mutant midbrain at E12.5.

## Acknowledgements

We thank Outi Kostia and Eija Koivunen for expert technical assistance. We thank Wolfgang Wurst for the *En1^Cre^* and Lori Sussel for the *Nkx2-2^Cre^* mice. We acknowledge the DNA Microarray Centre at Turku Centre for Biotechnology and Matti Kankainen and Daniel Borshagovski for the help in gene expression profiling and data analyses.

## Author contributions

J.P., K.A., M.S. and T.S. conceived and supervised the project. K.A. performed the microarray experiments. K.A., L.L. and P.Si. validated the microarray results and analyzed *Skor2* and *Nkx2-2* expression. Y.O. generated the *Skor2^GFP^* mice. A.K. analyzed the *Skor2* and *Nkx2-2* mutant mice, A.K., P.Si., A.M. and T.A.-A. performed the retrograde tracing experiments. Z.L. and A.K. designed and performed REM sleep deprivation experiments. S.M. designed and supervised, and S.M., P.Se. and P.Si. performed the electrophysiology measurements and recorded cell morphology analyses. S.K. performed the statistical testing and data normalization for the REM-sleep deprivation experiments and morphology analyses results. All the authors contributed to writing of the manuscript.

## Legends for Supplementary Tables and Figures

**Supplementary Table S1.** Downregulated genes in the E12.5 *Gata2^cko^* midbrain.

**Supplementary Table S2.** Complete list of down- and upregulated genes in E12.5 *Gata2^cko^* dorsal and ventral midbrain, identified in the microarray data analyses.

**Supplementary Table S3.** The gene ontology (GO) terms enriched among the genes downregulated in the *Gata2^cko^* midbrain. Online DAVID tool was used for the enrichment analysis.

**Supplementary Table S4.**
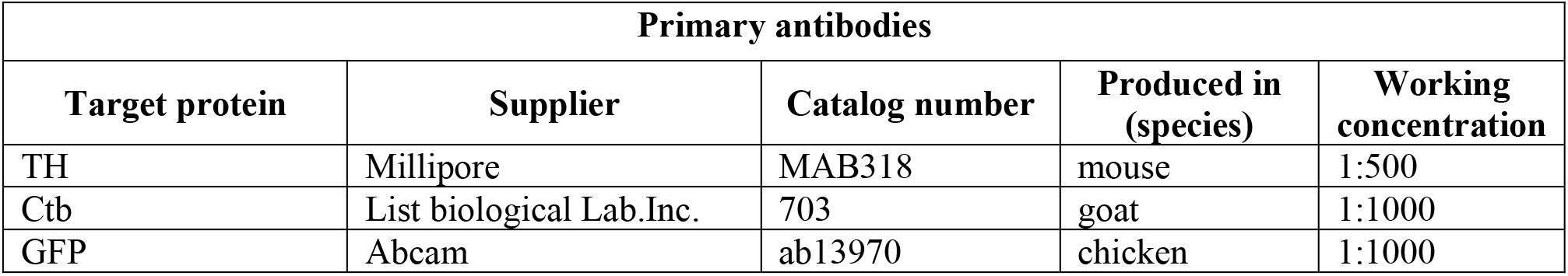
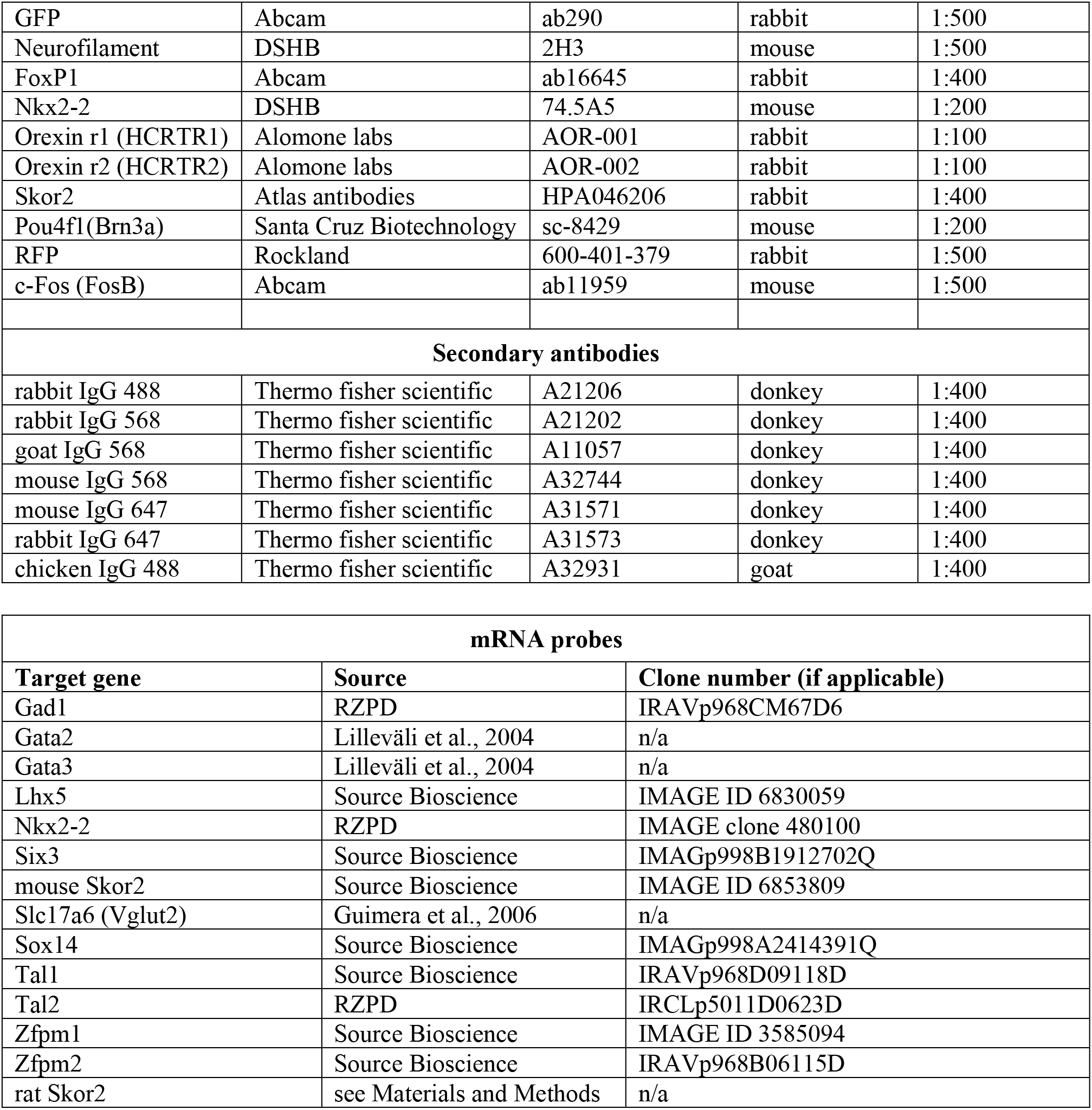
Antibodies and mRNA ISH probes used in the study.

**Supplementary Table S5.** The group identity, experiment ID, number of *Skor2*^+^ cells, number of c-Fos^+^ cells, the percent of c-Fos labelled *Skor2*^+^ cells and the transformed z-scores for each animal in the REM sleep deprivation assay.

**Supplementary Figure S1.**
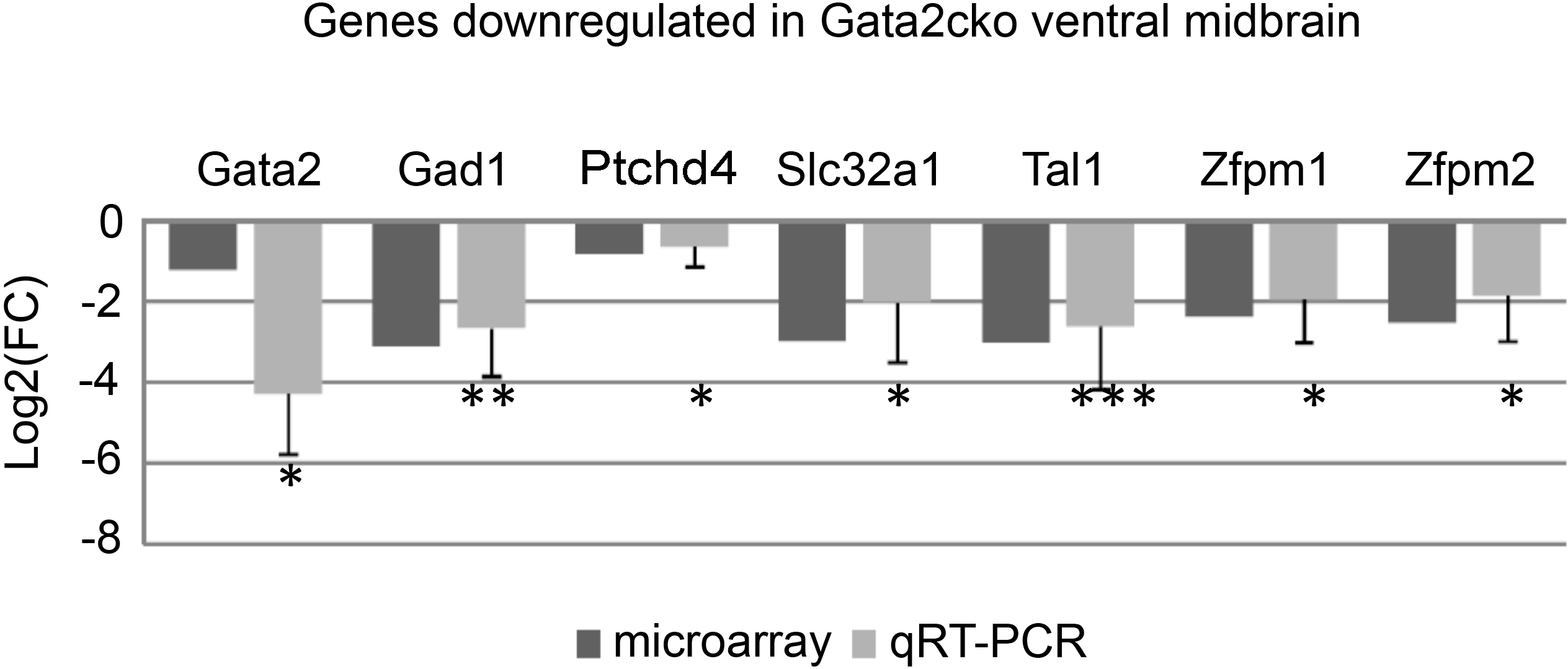
Comparison of the fold change indicated by the microarray data analysis and quantitative RT-PCR. FC, fold change. For a selection of predicted Gata2-regulated genes, qRT-PCR assay was performed. After RT-PCR, fold change was calculated using the Ct values detected in the *Gata2^cko^* and control (*En1^Cre^; Gata2^flox/wt^*) cDNA. Samples were normalized against *Actb* expression level in the same sample. The qRT-PCR was performed with 4 replicate cDNA samples. The statistical significance of the fold change in normalized expression levels is indicated. *** P<0.001, ** P<0.01, * P<0.05. The small fold-change of *Gata2* gene expression in the microarray is likely due to the fact that the microarray probe detects a truncated non-functional *Gata2* transcript.

**Supplementary Figure S2.**
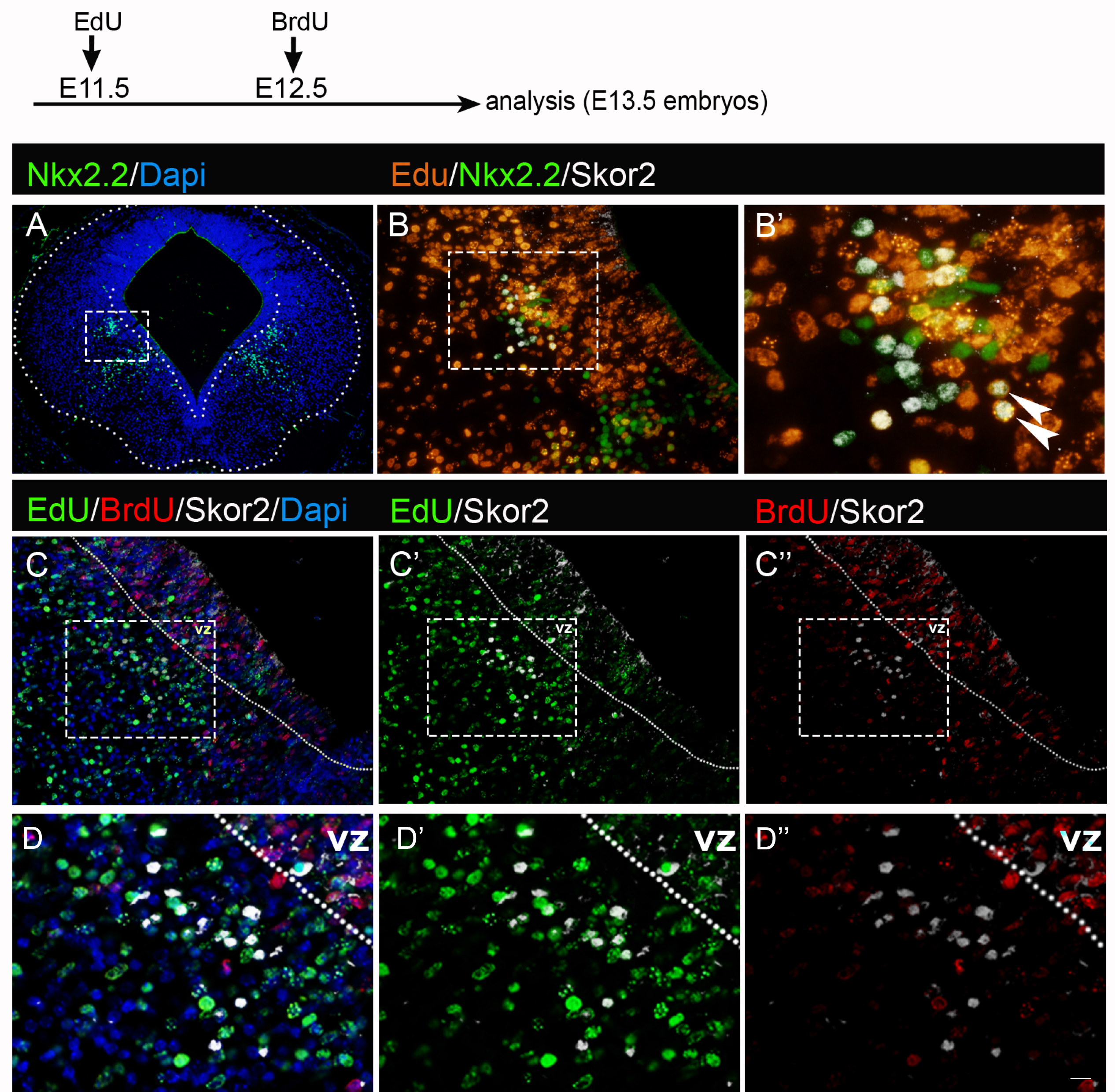
*Skor2*^+^ neuron birth-dating by BrdU and EdU labelling. Pregnant females received one dose of EdU at E11.5 and one dose of BrdU at E12.5 of pregnancy. Embryos were dissected at E13.5 and analyzed for the label. A, Nkx2-2 IHC on a coronal section of the E13.5 mouse midbrain. B-B’, Coexpression of Nkx2-2, Skor2 and EdU in E13.5 mouse midbrain. B’, shows a close-up of Skor2^+^ cells (dashed box in B). Arrowheads indicate triple-labelled cells. C-C’’, Labelling for EdU and BrdU in Skor2^+^ cells. EdU is detected in most of the Skor2^+^ cells (C’). BrdU is not detected in the Skor2^+^ cells (C’’). Close-ups of indicated regions are shown in D-D’’. vz, ventricular zone.

**Supplementary Figure S3.**
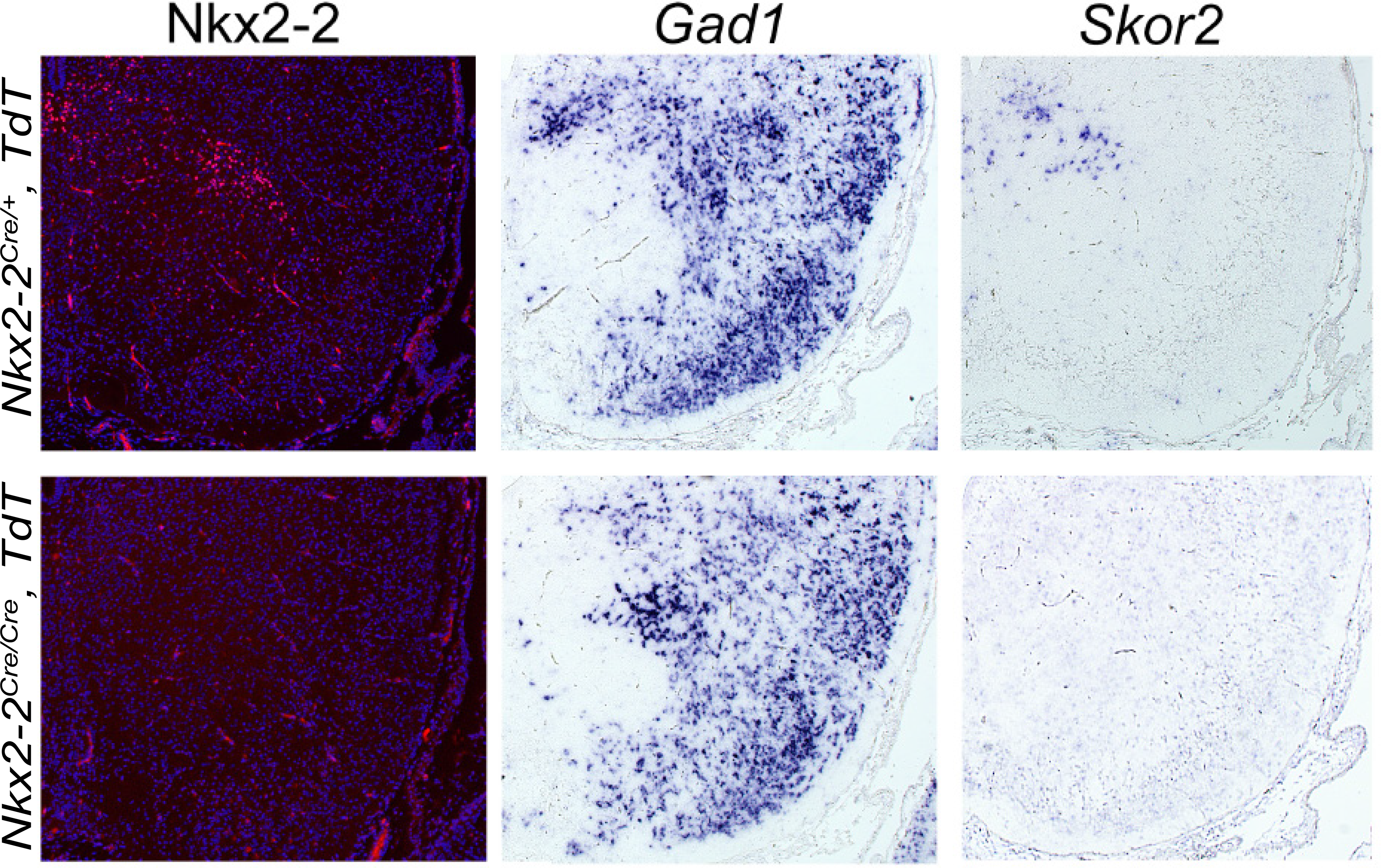
Skor2, but not Gad1 expression is lost in the Nkx2-2 mutant mouse. Coronal sections of E18.5 mouse midbrain analysed for the expression of Nkx2-2 (IHC), *Gad1* and *Skor2* (ISH), in *Ctrl* (*Nkx2-2^Cre/wt^, TdT*) and *Nkx2-2^null^* mutant (*Nkx2-2C^re/Cre^, TdT)* mice.

**Supplementary Figure S4.**
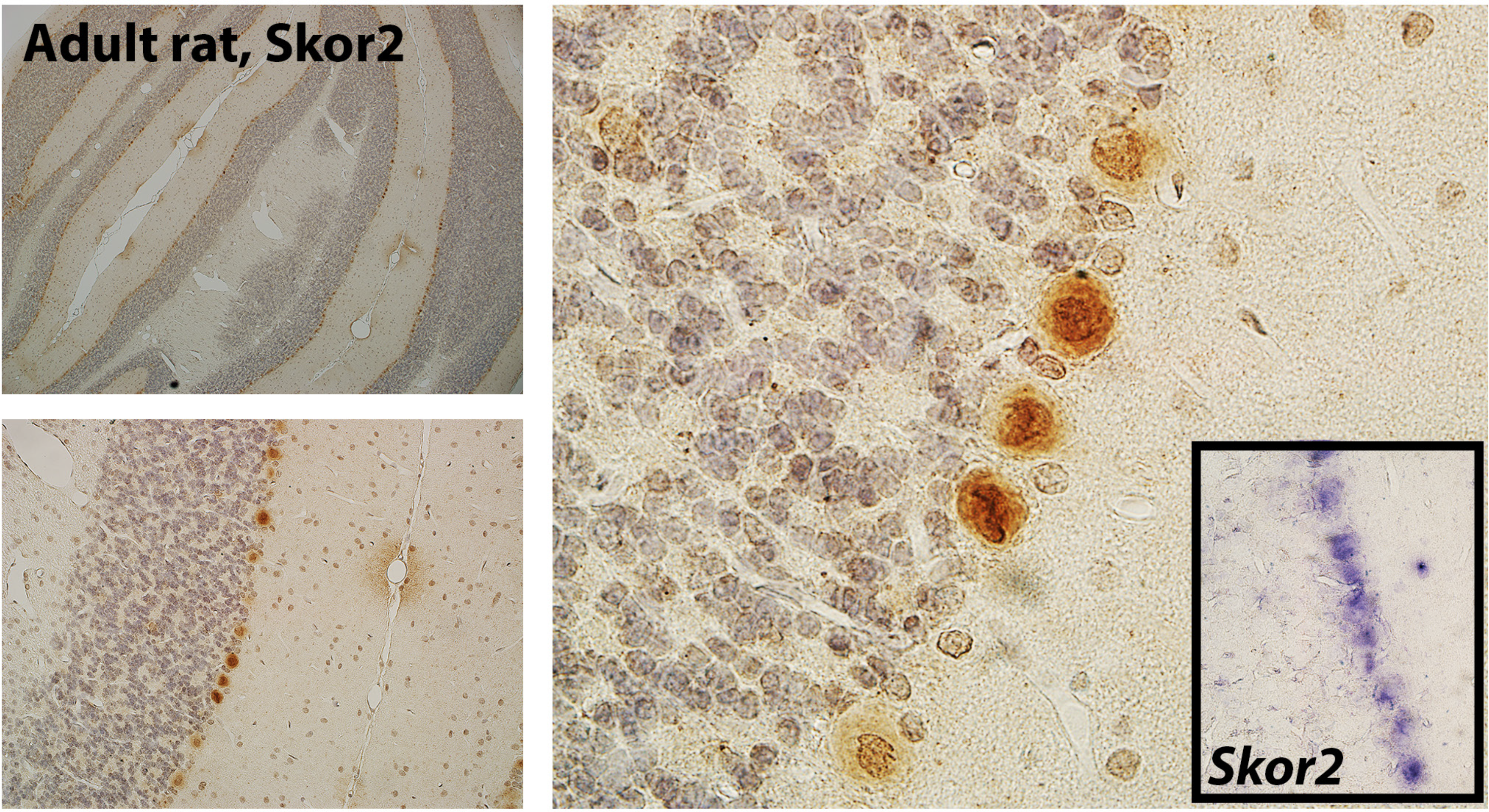
Skor2 expression in the adult rat cerebellum. IHC and ISH on coronal sections. Skor2 protein was detected in the Purkinje cell layer in the rat cerebellum (orange, IHC). The inset shows Skor2 mRNA expression in the Purkinje cell layer (violet, ISH).

**Supplementary Figure S5.**
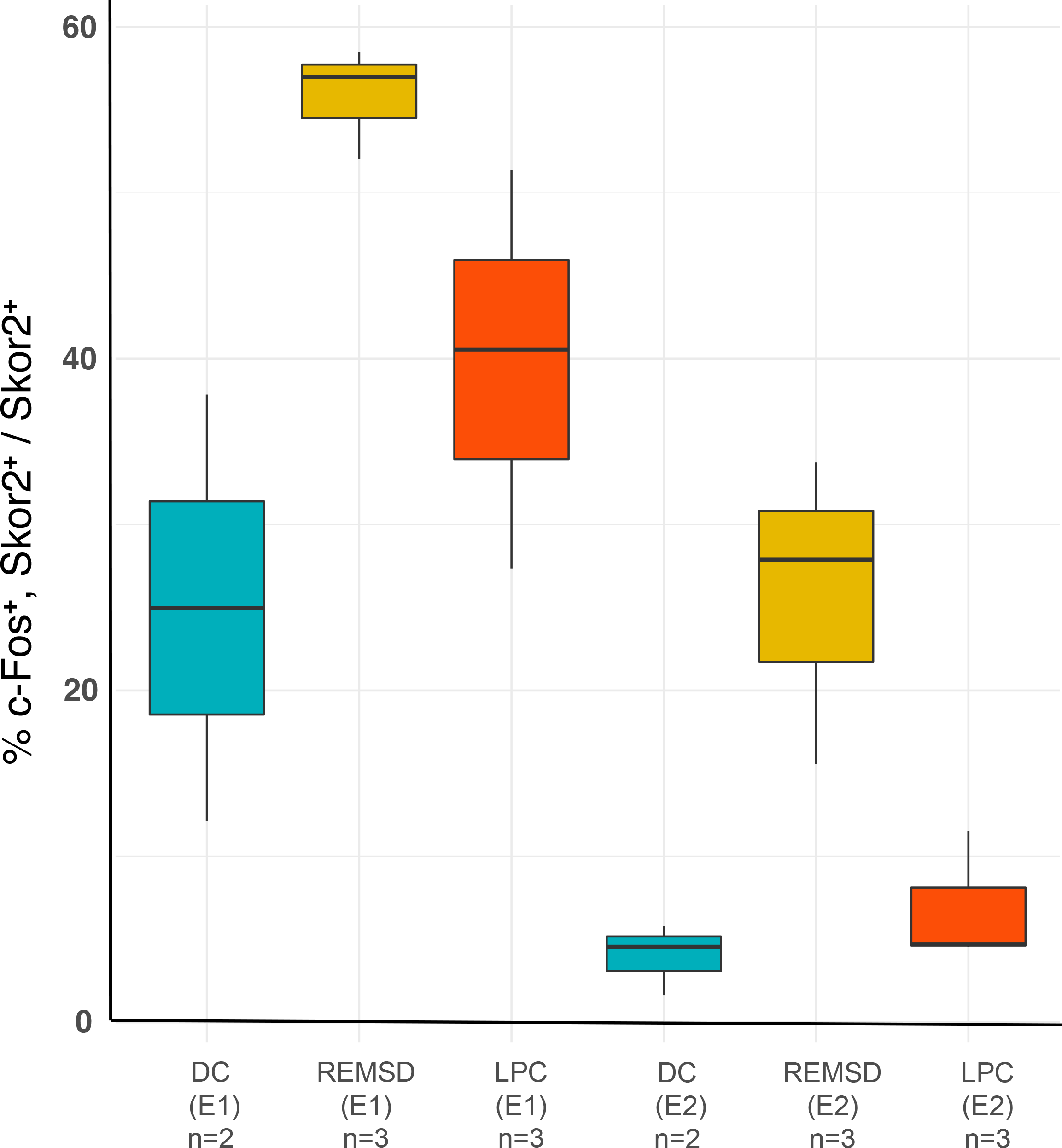
c-Fos staining efficiency in REM sleep deprivation experiments, before normalisation. Data is presented as percent of c-Fos-positive cells from the dMRF/vlPAG Skor2^+^ cells. Experiment groups (DC, REMSD, LPC) and experiment number (E1, E2) is indicated. Due to staining efficiency or other technical variation, the proportion of c-Fos labelled cells differs between the individual experiments (E1 and E2). The difference is systematic, as the experiment groups (DC, REMSD, LPC) show the same trend in labelled cell proportion. The number of animals (n) in each experiment and treatment group is indicated.

**Supplementary Figure S6.**
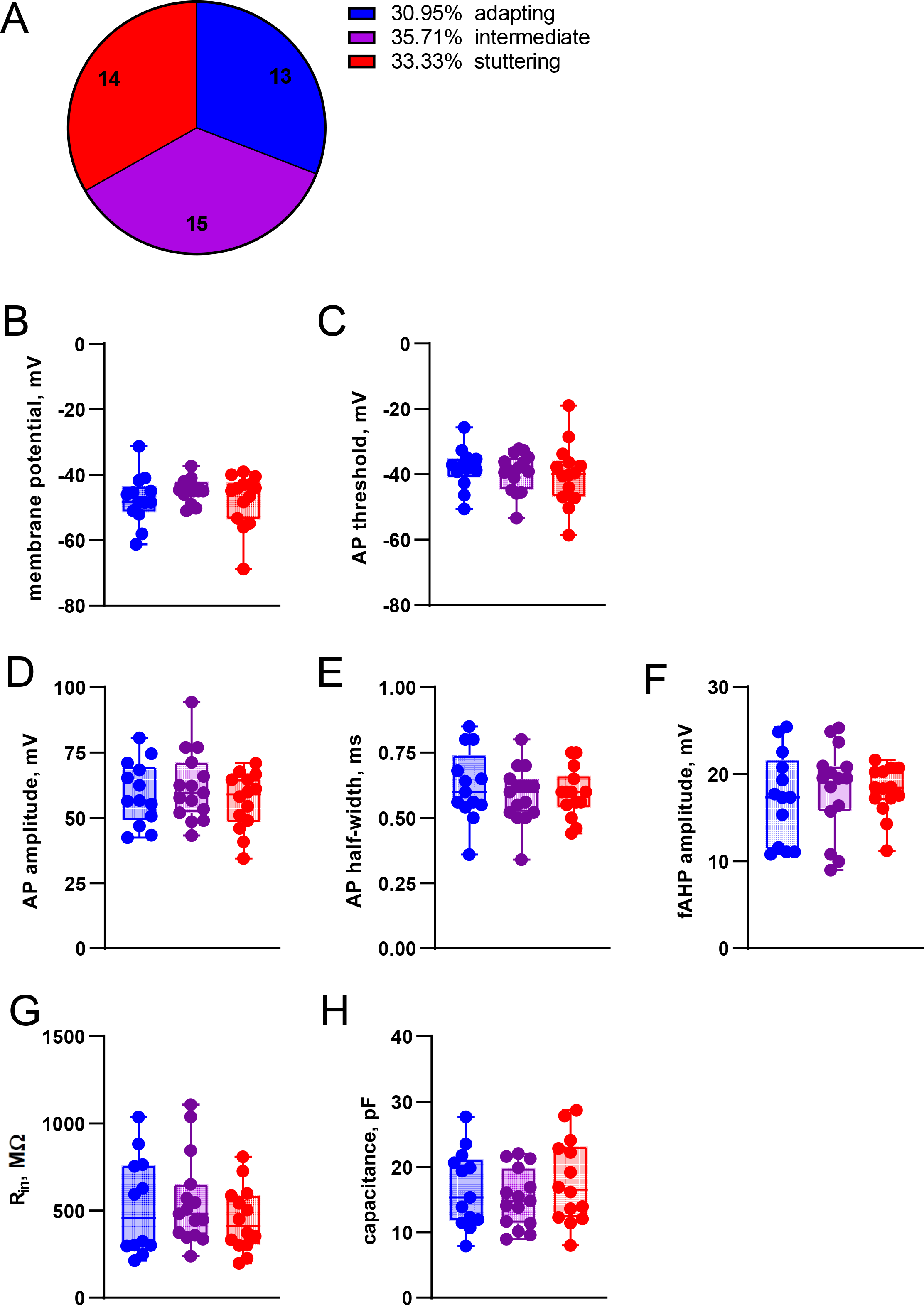
Classification of the *Skor2*^+^ neurons by the firing patterns. A, The proportion of cells by the assigned firing pattern class. B-H, Comparison of the excitability parameters measured in adapting, intermediate and stuttering Skor2^+^ neurons by electrophysiology. Boxplots show median, 25 and 75 percentiles with whiskers showing minimum and maximum values. RMP, resting membrane potential; AP, action potential; mAHP and fAHP, medium and fast after-hyperpolarizing potential; Rin, input resistance.

**Supplementary Figure S7.**
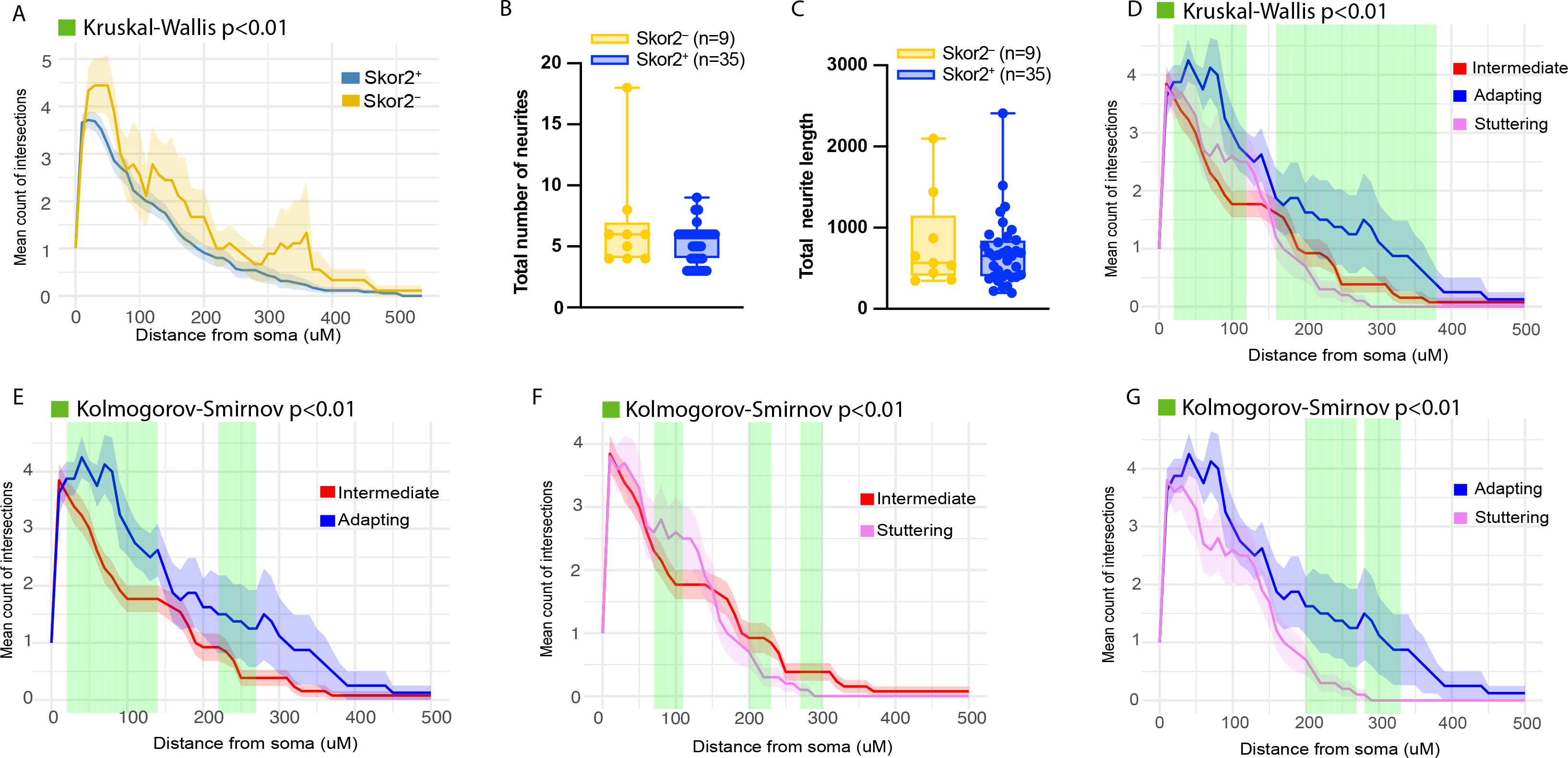
Comparison of morphological measurements of the *Skor2*+ and *Skor2*-neurons and the *Skor2*+ neuron subclasses. A, Average number of the intersections (+/- SEM) in Sholl analysis of *Skor2*^+^ (n= 35) and *Skor2*^-^ (n= 9) cells. The distance intervals where significant change (Kruskal-Walls test p<0.01) in the number of intersections was observed is highlighted in green. Calculations were conducted with moving window of 50 μm. B-C, Comparison of the neurite number (B) and length in pixels (C) between the of *Skor2*^+^ (n= 35) and *Skor2*^-^ (n= 9) cells. D-G, Average number of the intersections (+/- SEM) in Sholl analysis of *Skor2*+ neuron subclasses. The comparative analysis of all three cell classes (D), as well as the pairwise comparisons are shown. The distance intervals where significant change in the number of intersections was observed is highlighted in green. Kruskal-Wallis test of multiple comparisons was used in the comparison of three groups, Kolmogorov-Smirnov test for the pairwise comparison. Significance threshold in both tests was p<0.01. Calculations were conducted with moving window of 50 μm.

